# Genome evolution and convergent innovation in the carnivorous plant *Sarracenia purpurea*

**DOI:** 10.64898/2025.12.26.696377

**Authors:** Victor A. Albert, Jonathan Kirshner, Christopher Page, Nicholas Pratt, Johanna Merkel, Michaela Richter, Matthias Freund, Kenji Fukushima, Charlotte Lindqvist

## Abstract

The repeated evolution of certain complex traits raises a fundamental question of how genomes generate ecological novelty while preserving developmental stability. Carnivorous pitcher plants, which modify a core organ of plant performance, the leaf, exemplify this challenge, yet the genomic basis of their convergent evolution has not been resolved. We present a chromosome-scale genome assembly for *Sarracenia purpurea* and analyze it alongside eight additional angiosperms spanning its parent clade Ericales and several other carnivorous lineages. The genome reveals extensive syntenic duplicate blocks arising from ancient polyploidy events, with diffuse, alternating dominant and recessive segments interleaved along chromosomes. Dominant regions are biased toward retained copies of dosage-sensitive regulatory genes, including *AGO1*, *BRX*, *GATA11*, *ETC1* and *RCD1*. These genes highlight a conserved regulatory scaffold associated with leaf morphogenesis, including epidermal differentiation, auxin-mediated patterning, and redox-integrated coordination. By contrast, tandem gene duplications preferentially accumulate in structurally labile genomic regions and constitute a complementary, rapidly evolving component of the genome, enriched for ecological effector functions, including detoxification, glutathione-mediated redox buffering, antifungal pathways, and cuticle modification activities. Comparative union-based functional analyses across four carnivorous taxa reveal convergent recruitment of oxidative, transport, microbial-interaction, and cell-wall processes during independent trap evolution. Transcriptomic data confirm consistent activation of these pathways in pitchers. These findings demonstrate that complex traits arise through a genome-wide partitioning between polyploidy-derived, conserved developmental regulation and tandem-driven ecological specialization, here partitioning leaf architectural control from rapidly evolving functions associated with pitfall-based prey capture.

## 1. Introduction

Carnivorous plants have long attracted scientific attention because they display remarkable evolutionary innovations in leaf architecture, nutrient acquisition, and ecological strategy^1,2^. Among these lineages, the North American pitcher plant *Sarracenia purpurea* stands out for its distinctive fluid-filled traps that collect, retain, and digest arthropod prey^3^ (**Figure 1**). The traps support a complex digestive ecosystem in which microbes, small invertebrates (such as larval dipterans), and decomposing prey interact in a structured trophic network^4–12^. Prey decomposition and nutrient release depend heavily on microbial metabolism, with relatively limited direct enzymatic contribution from the plant, particularly when compared with other pitcher plant genera such as *Nepenthes*, which produces enzyme-rich secretions^13–15^. Although pitcher morphology is a classic example of convergent evolution among *Sarracenia* (the North American pitcher plant), *Cephalotus* (the Australian pitcher plant), and *Nepenthes* (the Southeast Asian pitcher plant)^16^, the genomic mechanisms by which these lineages independently reprogram leaf development and implement distinct digestive strategy remain poorly understood.

**Figure 1.**
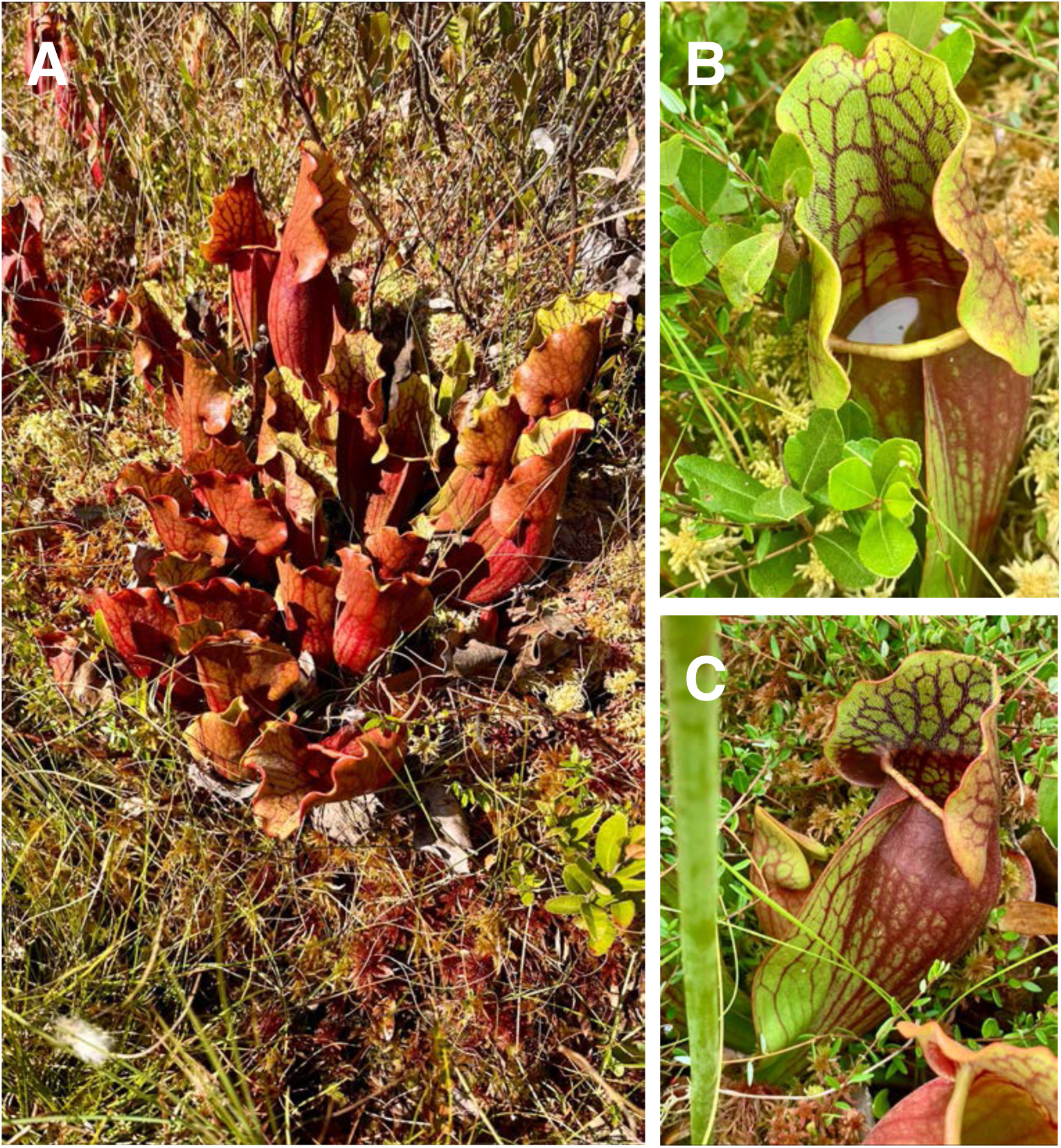
*Sarracenia purpurea*. **(A)** Whole plant forming a clump of pitchers, **(B)** a single pitcher showing the lid, peristome around the trap opening, and fluid-filled pitcher tube, and **(C)** a side view of a pitcher highlighting the conspicuous wing. Photos by V.A.A **(A)**, from Allenburg Bog Audubon Nature Reserve, New York; **(B)** and **(C)**, from Mendon Ponds Park, New York).

*Sarracenia purpurea* therefore offers a unique opportunity to study how a plant can reconfigure leaf function to support a stable micro-ecosystem that performs much of the digestive work. In the absence of a chromosome-scale genome, however, it has not been possible to evaluate how genome structure and gene duplication contribute to pitcher development and digestive ecology, or how these features relate to broader patterns of pitcher plant genome evolution.

Despite considerable phenotypic and molecular developmental^17^, environmental (e.g., ^18^), and marker-based phylogenetic research (e.g.,^19,20^), the genomic foundations of pitcher-leaf development and digestive biology in *Sarracenia* have not yet been examined at chromosome scale. Earlier genomic studies of carnivorous plants have focused on systems such as *Utricularia*^21^, *Cephalotus*^22^, and *Nepenthes*^23^, where genome reduction (in the former) or convergent trapping strategy (in the latter two) has motivated prior genomic research. Without a resolved genome for *Sarracenia*, the ways in which distinct modes of gene duplication shaped pitcher-plant functional repertoires, and how duplicated genes were differentially retained or lost, have remained unclear, rendering comparative research on pitcher morphology and digestive function incomplete. While the incompleteness of the *Cephalotus* assembly^22^ has limited productive comparison with *Nepenthes*, even in the absence of a fully resolved *Cephalotus* genome, a chromosome-scale *Sarracenia* assembly enables informative pairwise and multi-lineage inference.

*Sarracenia* belongs to the Ericales, a clade believed to have originated from an ancient allopolyploidy event^24^. Ericales includes both carnivorous and many familiar noncarnivorous lineages, from woody trees and shrubs to herbaceous species, and it encompasses a wide diversity of ecological strategies^25^. A chromosome-scale *Sarracenia* genome, aligned against multiple Ericales relatives, therefore allows two complementary questions to be addressed. First, which structural features and duplicated blocks are shared and thus represent the ancestral genomic framework within which *Sarracenia* evolved carnivory. Second, which rearrangements, fractionation patterns, and regional gene family expansions are specific to *Sarracenia* and may reflect lineage-specific redeployment of duplicated genes during pitcher evolution.

Understanding the evolution of *Sarracenia* requires an integrative framework that combines structural genomics, syntenic comparisons, synonymous substitution rate (Ks)-based event dating interpreted in light of mixture components recovered for *Sarracenia* and other Ericales, and gene functional enrichment analyses. Synteny identifies conserved blocks inherited from the ancestral core eudicot (and the *gamma* allohexaploidy that marks it^26,27^) and reveals how these blocks were rearranged following whole-genome duplication. Ks distributions provide information about the relative timing of duplication events, both whole-genome and small scale (tandem), through correspondence with background mutation rates. Cross-comparisons with taxa such as *Vitis vinifera* (grapevine, an early-branching rosid) help anchor these events on a relative timescale because of its conservative Ks behavior and modest structural evolutionary rate^28–30^. Subgenome reconstruction clarifies how differential retention versus loss of duplicate genes shaped dominant and recessive genomic compartments, a post-polyploid feature that may influence the stability of regulatory networks while permitting divergence of functionally flexible gene families^31–35^. In combination, these approaches make it possible to reconstruct the genomic scaffolding that supports *Sarracenia* pitcher-leaf development and digestive function.

Carnivorous plants have evolved their traps repeatedly, and this repetition creates a natural comparative context^2^. By placing *Sarracenia* alongside *Nepenthes gracilis* (Caryophyllales), *Pinguicula gigantea* and *Utricularia gibba* (Lamiales), and by embedding all of them in a phylogenetic framework that includes both closely and more distantly related non-carnivores, it becomes possible to distinguish convergent solutions from lineage-specific innovations and background traits inherited from non-carnivorous ancestors. A nine-taxon Venn-bin analysis, which includes *Sarracenia*, *Nepenthes*, *Pinguicula*, *Utricularia*, *Vitis*, *Arabidopsis thaliana* (Brassicales, and our annotation reference), *Actinidia chinensis* (Ericales), *Mimulus guttatus* (Lamiales), and *Beta vulgaris* (Caryophyllales), separates syntenically retained regulatory genes from tandem-duplicate driven functional innovation and allows functional categories to be traced across shared, convergent, and lineage-specific contexts. Because *Actinidia* is the only Ericales representative in this dataset, it also serves as a filter for removing functional enrichments that reflect general Ericales biology rather than carnivory-associated innovation.

Here, we present a high-quality chromosome-level genome assembly of *Sarracenia purpurea*, a detailed gene and repeat annotation, an analysis of self-synteny patterns that reveals the imprint of ancient whole-genome duplication and subsequent rearrangement, a multi-species Ericales anchoring framework that situates *Sarracenia* genome structure within its broader clade, a reconstruction of *Sarracenia* subgenomes using *Vitis* as an outgroup, and an evolutionary-functional analysis using intersecting duplicate-gene and gene ontology (GO) datasets as foregrounds for enrichment tests. We also explore pitcher-specific gene expression data for correlates of trap function. These analyses clarify how distinct modes of gene duplication partition developmental regulation from ecological function in a carnivorous leaf, providing a genomic perspective on the repeated evolution of pitcher plants.

## 2. Results and Discussion

### 2.1 Genome Assembly, Annotation, and Ericales Anchoring Context

#### 2.1.1 Chromosome-Level Assembly of *Sarracenia purpurea*

The *Sarracenia purpurea* genome was assembled from high-coverage Oxford Nanopore Technology long reads^36^ and reference-scaffolded against another *Sarracenia* genus assembly into chromosome-scale sequences. The total length across 4,071 contigs was 3,523,392,364 bp, with an N50 of 2,875,369 bp (**Table 1**). The final scaffolded assembly of 3,523,623,004 bp comprises 13 chromosome-length scaffolds (pseudochromosomes), consistent with the expected haploid chromosome number. Assembly continuity is high, with a scaffold N50 of 241,630,572 bp, and BUSCO^37^ analysis shows 95.5% completeness for the genome and 84.1% completeness for the annotated gene set (see more detail on annotation in section 2.1.2). Large tracts of reduced gene density mark presumed pericentromeric and centromeric regions (**Figure 2**). These features provide a coherent chromosomal framework that supports subsequent synteny analyses, reconstruction of ancient homoeologous segments, and subgenome inference.

**Figure 2.**
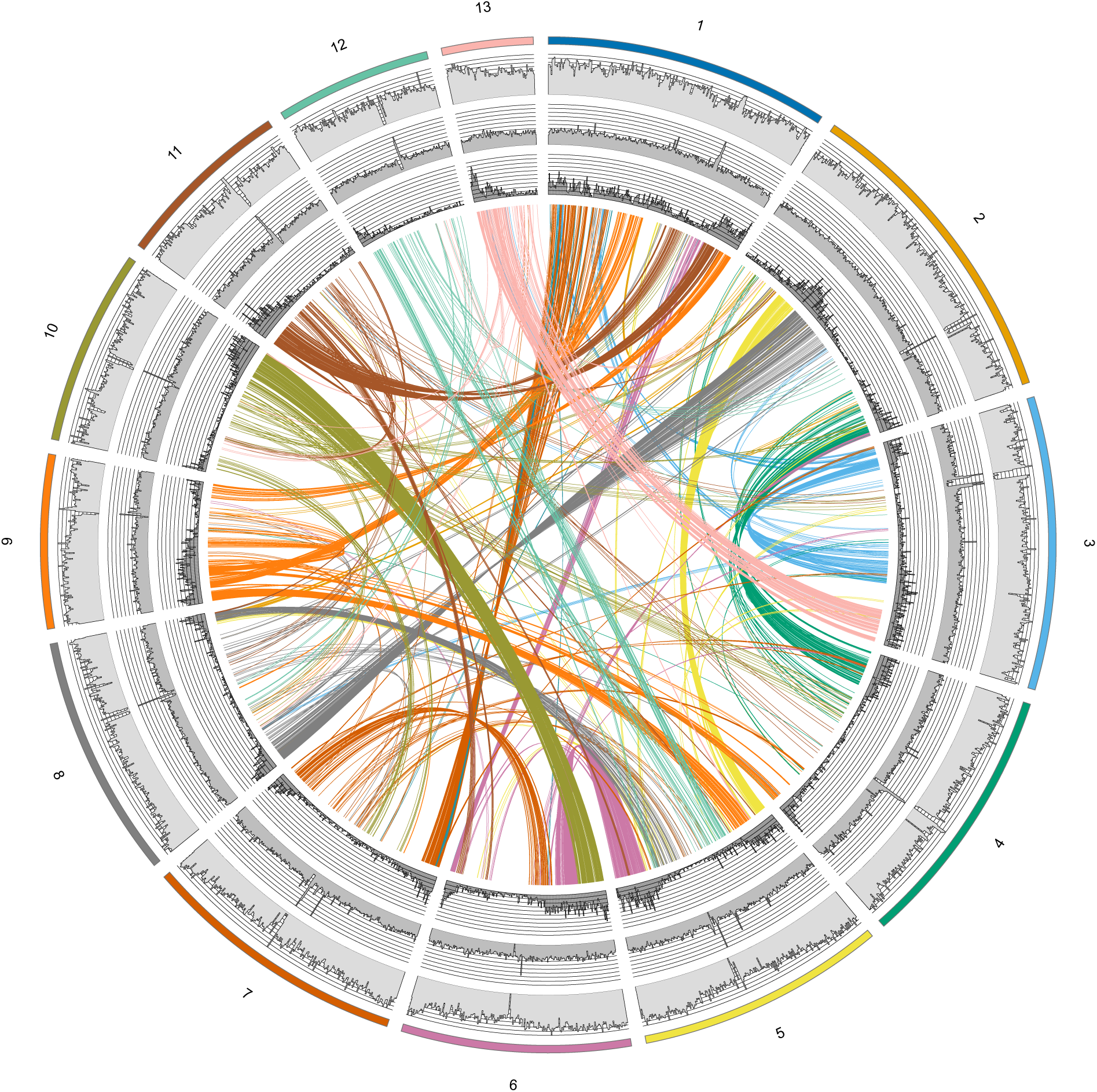
Chromosome-wide genomic feature profiles in *Sarracenia purpurea*. Normalized tracks along each of the 13 chromosomes show gene density, LTR retrotransposon abundance, total repeat content, and locations of self-syntenic blocks inferred from duplicated anchor pairs. Gene density is displayed as a continuous track, with higher values in darker or taller segments. Separate color-coded tracks distinguish total LTR content and major LTR subfamilies (for example, Gypsy and Copia), and a further track marks overall repetitive content. Self-syntenic blocks are drawn as colored segments spanning collinear runs of duplicated genes, the connecting arcs restricted to blocks with log10(Ks) < 1. Chromosome labels and physical coordinate scales along the outer ring allow direct comparison of gene-rich, TE-rich, and duplication-rich regions across the genome.

**Table 1.** Global assembly and gene space statistics for the *Sarracenia purpurea* reference genome. Columns include total assembled length, number of contigs and scaffolds, contig N50 and L50, scaffold N50 and L50, largest contig and scaffold sizes, GC content, and counts of contigs and scaffolds above specified size thresholds. BUSCO statistics are provided for the both contig- and scaffold-level assemblies, as well as for the genome annotation. These metrics confirm a high-quality contig assembly, the completeness of the gene space, and confirm chromosome-scale scaffolding for the 13 pseudochromosomes.

| METRICS: GENOME | FILTERED CONTIG ASSEMBLY | RAGTAG SCAFFOLDED |
| --- | --- | --- |
| COMPLETE BUSCOS (C) | 2210 (95.0%) | 2223 (95.5%) |
| COMPLETE AND SINGLE-COPY BUSCOS (S) | 2007 (86.3%) | 2036 (87.5%) |
| COMPLETE AND DUPLICATED BUSCOS (D) | 203 (8.7%) | 187 (8.0%) |
| FRAGMENTED BUSCOS (F) | 35 (1.5%) | 26 (1.1%) |
| MISSING BUSCOS (M) | 81 (3.5%) | 77 (3.4%) |
| TOTAL BUSCOS SEARCHED | 2326 | 2326 |
| # CONTIGS OR SCAFFOLDS | 4071 | 1884 |
| LARGEST CONTIGS OR SCAFFOLDS | 45271496 | 321174296 |
| TOTAL LENGTH | 3523392364 | 3523623004 |
| GC (%) | 39.04 | 39.04 |
| N50 | 2875369 | 241630572 |
| N90 | 591864 | 7662836 |
| L50 | 325 | 7 |
| L90 | 1334 | 26 |
| N'S PER 100 KBP | 0 | 6.22 |
| **METRICS: ANNOTATION** |  |  |
| COMPLETE BUSCOS (C) |  | 1956 (84.1%) |
| COMPLETE AND SINGLE-COPY BUSCOS (S) |  | 1763 (75.8%) |
| COMPLETE AND DUPLICATED BUSCOS (D) |  | 193 (8.3%) |
| FRAGMENTED BUSCOS (F) |  | 173 (7.4%) |
| MISSING BUSCOS (M) |  | 197 (8.5%) |
| TOTAL BUSCOS SEARCHED |  | 2326 |

#### 2.1.2 Annotation and Gene Space Characterization

The annotated gene set shows characteristics typical of a core eudicot. Transcript-supported models from trap, flowers and root tissues helped ensure that both broad functions, including trap-associated genes, are retained in the annotation. The filtered, nonredundant annotation includes 28,835 protein-coding loci, each represented by a single transcript model. Mean exon length is ∼263 bp, annotated genes contain an average of ∼4-5 exons, and ∼25% of loci are single-exon genes (**Table S1**).

Gene density varies substantially across the genome. Using 1-Mb windows across chromosomes larger than 5 Mb, gene density ranges from 0 genes/Mb in repeat-rich interior regions to a maximum of 42 genes/Mb in more euchromatic segments, reflecting strong spatial heterogeneity in gene distribution **(Figure 3)**. At the genome-wide scale, the distribution of 1-Mb window gene densities is strongly right-skewed, with many low-gene windows and a long tail of gene-dense windows **(Figure 4A).**

**Figure 3.**
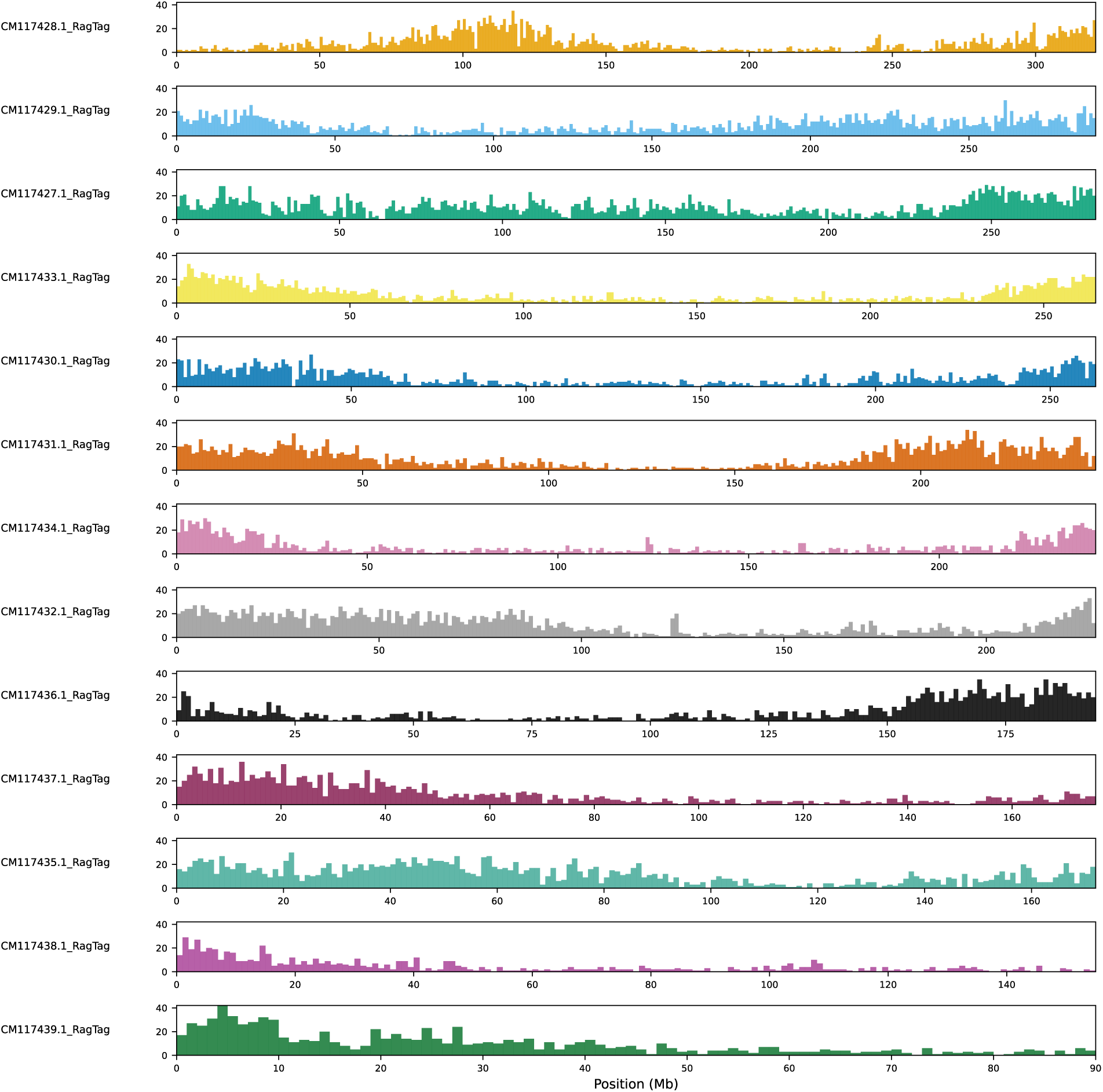
Chromosome-scale gene-density variation across the *Sarracenia purpurea* pseudochromosomes. Gene density was quantified in non-overlapping 1-Mb windows across the 13 RagTag pseudochromosomes (CM*). Bars show the number of annotated genes per window (gene midpoints assigned to windows). The tracks highlight strong spatial heterogeneity, including extended gene-poor regions consistent with repeat-rich compartments and localized peaks of elevated gene density in more gene-rich segments.

**Figure 4.**
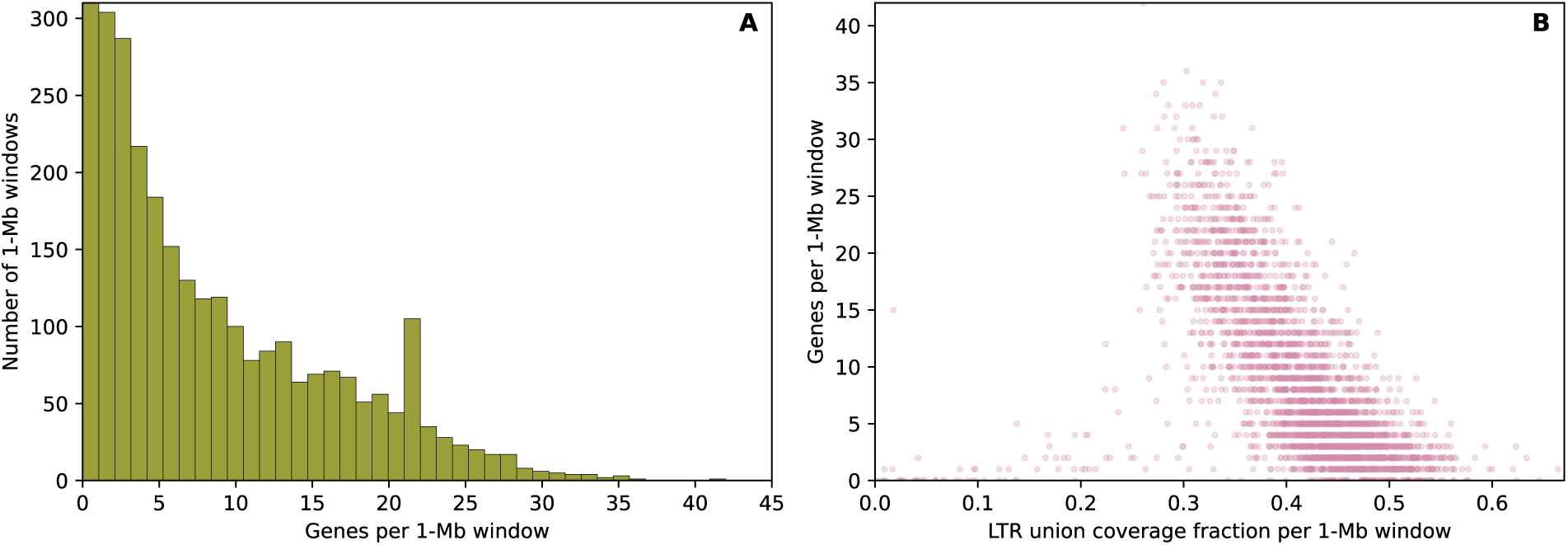
Genome-wide distribution of 1-Mb gene density and its relationship to LTR coverage. **(A)** Histogram of genes per 1-Mb window across the 13 pseudochromosomes, showing a strongly skewed distribution with many low-gene windows and a long tail of gene-dense windows. **(B)** Scatterplot relating gene density (genes per 1-Mb window) to LTR union coverage fraction in the same windows, showing that LTR-rich windows tend to be gene-poor, consistent with megabase-scale compartmentalization of gene-rich and repeat-rich regions.

Transposable elements and other interspersed repeats constitute the majority of the *Sarracenia purpurea* genome (**Table S2**). RepeatMasker annotation^38^ indicates that ∼88% of the assembly is masked, of which ∼86.6% corresponds to interspersed repeats. At the level of TE class composition, LTR (long terminal repeat) retrotransposons dominate, comprising ∼71.8% of the genome, with Copia elements representing ∼33.4% and Gypsy elements ∼20.5%. DNA transposons account for an additional ∼1.1%, while the remaining repeat fraction is distributed among unclassified elements (∼12.4%) and other low-abundance families. Together, these proportions indicate that the *Sarracenia* genome has been shaped primarily by extensive LTR retrotransposon accumulation at the genome-wide scale.

This overall dominance, however, masks substantial variation across individual pseudochromosomes. When TE coverage is quantified at the scaffold level using union RepeatMasker coverage, overall TE occupancy (all classes combined) varies from ∼84.3-90.6% across the 13 pseudochromosomes (Δ ∼6.3 percentage points; coefficient of variation ∼2%), while LTR-tagged coverage specifically ranges from ∼38.9-45.3% (Δ ∼6.3 percentage points; CV ∼4%). Thus, although TE distributions vary among scaffolds, LTR retrotransposons remain the principal contributors to repeat content throughout the genome. This combination of genome-wide LTR dominance and chromosome-scale heterogeneity is consistent with repeated cycles of LTR expansion, removal, and rearrangement contributing to genome size variation, regional differences in gene density, and erosion of ancient homoeologous segments following any ancestral whole-genome duplications (WGDs). Consistent with this compartmentalization, 1-Mb windows with higher LTR coverage tend to be gene-poor (**Figure 4B**).

Divergence landscapes derived from RepeatMasker annotations provide an additional temporal perspective on repeat accumulation in *Sarracenia* (**Figure 5**). All RepeatMasker-annotated repeat fragments were binned by their reported percent divergence from consensus (integer bins from 0-50%), and base-pair coverage was summed per bin to generate bp-weighted divergence profiles. The resultant genome-wide repeat landscape is dominated by a mode centered near ∼7% divergence, whereas LTR-tagged repeats show a peak near ∼12% divergence. Because RepeatMasker divergence values are measured relative to consensus sequences and do not inherently provide absolute insertion times, the reported profiles in **Figure 5** are interpreted as reflecting relative patterns of repeat turnover within the *Sarracenia* genome. The separation between the genome-wide and LTR-restricted profiles indicates that LTR retrotransposons contribute disproportionately to a temporally structured component of repeat accumulation. These divergence landscapes complement chromosome-scale feature tracks (**Figures 2 and 3**) by showing that repeat proliferation has not been uniform through time and that LTR dynamics represent a major axis of genome-wide repeat remodeling.

**Figure 5.**
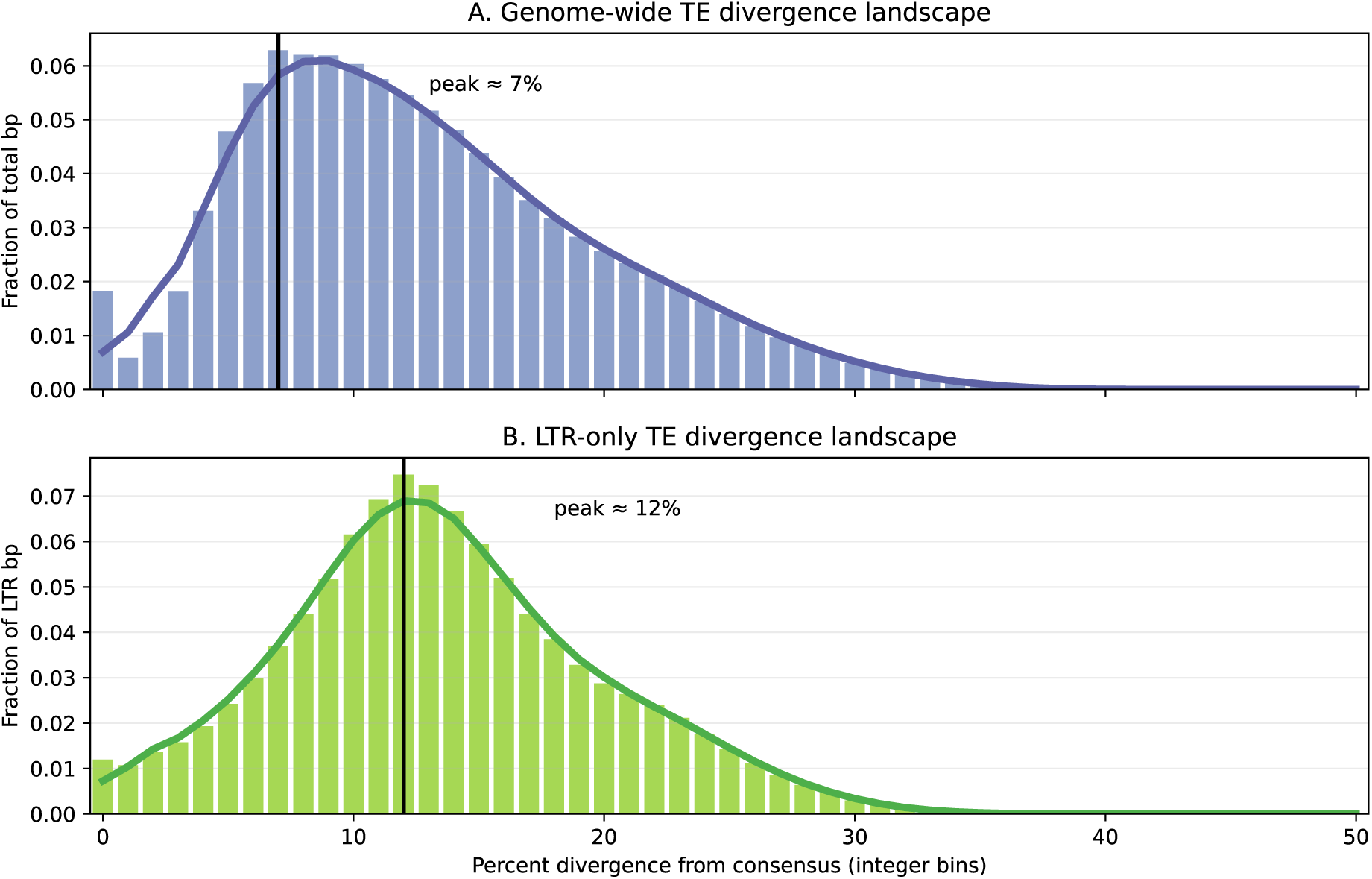
RepeatMasker divergence landscapes for *Sarracenia purpurea* repeats (genome-wide versus LTR-only). **(A)** Genome-wide repeat divergence landscape. **(B)** LTR-tagged repeat divergence landscape. For each RepeatMasker-annotated repeat fragment, the reported percent divergence from its consensus was binned into integer divergence bins (0-50%), and base-pair coverage was summed per bin to generate bp-weighted divergence profiles. Histograms show the fraction of total covered bp contributed by each divergence bin (panel-specific normalization), and the overlaid smoothed curves summarize the same distributions to highlight modal structure. Vertical reference lines mark the peak (modal) divergence bin for each panel.

#### 2.1.3 *Sarracenia* Self-Synteny and Structural Genomic Landscape

Self-synteny analysis (based on use of SynMap in CoGe^39–41^) reveals that the *Sarracenia purpurea* genome preserves large duplicated chromosomal segments traceable to an ancient WGD (**Figure 2**; cf. ^42^). Several chromosomes retain extended regions of near-collinear duplicated gene order, indicating conservative evolution in those regions since the WGD. Elsewhere, the synteny signal is broken into shorter and more fragmented blocks, likely reflecting a combination of gene loss, small-scale rearrangements, and local TE insertions. Consistent with this interpretation, genome-wide windowed analyses show a strong negative association between self-syntenic anchor density and repeat abundance, with megabase-scale windows lacking detectable syntenic anchors exhibiting substantially higher transposable-element density than anchor-rich windows (Spearman’s ρ ≈ −0.5; P ≪ 0.001), indicating that regions of syntenic erosion are embedded within repeat-rich genomic environments. Syntenic block Ks distributions^43,44^ show a prominent set of two components with major modal densities at ∼0.65 and ∼1.7 (**Figure 6**).

**Figure 6.**
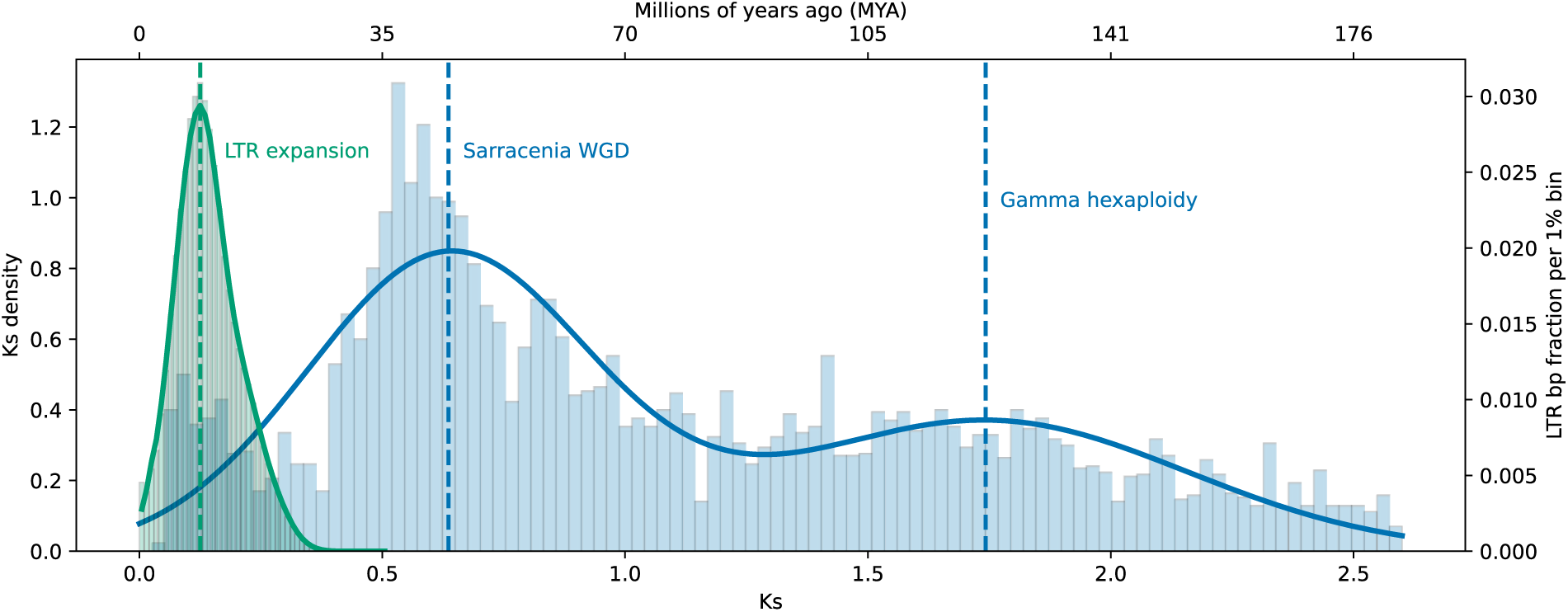
Temporal relationship between gene duplication and LTR retrotransposon expansion in *Sarracenia purpurea*. Genome-wide Ks distributions for self-syntenic duplicate gene pairs in *Sarracenia purpurea* are compared with RepeatMasker-derived divergence landscapes for LTR retrotransposons using a shared, calibrated time axis. Blue histogram and curve show the density of Ks values transformed to time, with dashed vertical lines indicating the means of the two dominant Ks components corresponding to a younger, lineage-specific duplication signal and a broader, older component associated with the ancient core-eudicot *gamma* hexaploidy. Green histogram and curve shows the fraction of LTR-associated base pairs per divergence bin, with the dashed vertical line marking the modal LTR expansion. Time scaling was calibrated by constraining the *gamma*-associated Ks component to ∼120-125 million years ago; resulting ages are therefore interpreted comparatively rather than as absolute insertion times. The temporal offset between the LTR expansion peak and both Ks components indicates that large-scale gene duplication and subsequent fractionation preceded the major episode of LTR proliferation in the *Sarracenia* genome.

To relate these duplication signals to the temporal structure of repeat accumulation, we compared the Ks distribution of self-syntenic duplicate pairs with RepeatMasker-based divergence landscapes for LTR retrotransposons using a common, calibrated time axis (**Figure 6**). This comparison reveals a clear separation between a young, lineage-specific LTR expansion and the two dominant duplication components inferred from Ks data. The LTR divergence profile is sharply peaked at low divergence, indicating a recent burst of LTR activity at ca. 10 Mya, whereas the Ks distribution resolves into a younger *Sarracenia*-specific duplication component at ca. 45-50 Mya and a broader, older component corresponding to the ancient core-eudicot *gamma* hexaploidy event at ca. 120-125 Mya (cf. ^45–47^). The lack of temporal overlap between the LTR expansion peak and either Ks component suggests that large-scale duplication and subsequent fractionation preceded the major episode of LTR proliferation, reinforcing the view that repeat accumulation has preferentially targeted regions already subject to syntenic erosion rather than driving whole-genome duplication itself.

#### 2.1.4 Multispecies Ericales Comparative Anchoring and Duplication Timing

To investigate the temporal structure of gene duplication within this framework, we examined Ks distributions for self-syntenic duplicate gene pairs. All Ericales taxa retain a broad, high-Ks component consistent with deeply diverged paralogs associated with the ancient core-eudicot *gamma* hexaploidy. In preliminary work, we found that several taxa also display prominent lower-Ks components reflecting more recent duplication activity, such as *Actinidia* and *Vaccinium*. The presence of these more recent whole-genome duplications obscures separation between *gamma*-derived and post-*gamma* duplication signals and complicates direct cross-taxon comparison of younger Ks modes. For this reason, although *Actinidia* and *Vaccinium* provide important structural anchoring context (see section 2.2), they were not used for comparative inference of post-*gamma* duplication timing.

Focusing instead on taxa with simpler duplication landscapes, *Sarracenia*, *Rhododendron*, and *Impatiens* reveals a clearer comparative signal. In addition to the shared *gamma* component, each of these genomes displays a distinct younger Ks mode. When calibrated independently using the *gamma* component within each species, these younger modes correspond to inferred ages of approximately 45 million years ago in *Sarracenia*, 58 million years ago in *Rhododendron*, and 59 million years ago in *Impatiens* (**Figure 7**).

**Figure 7.**
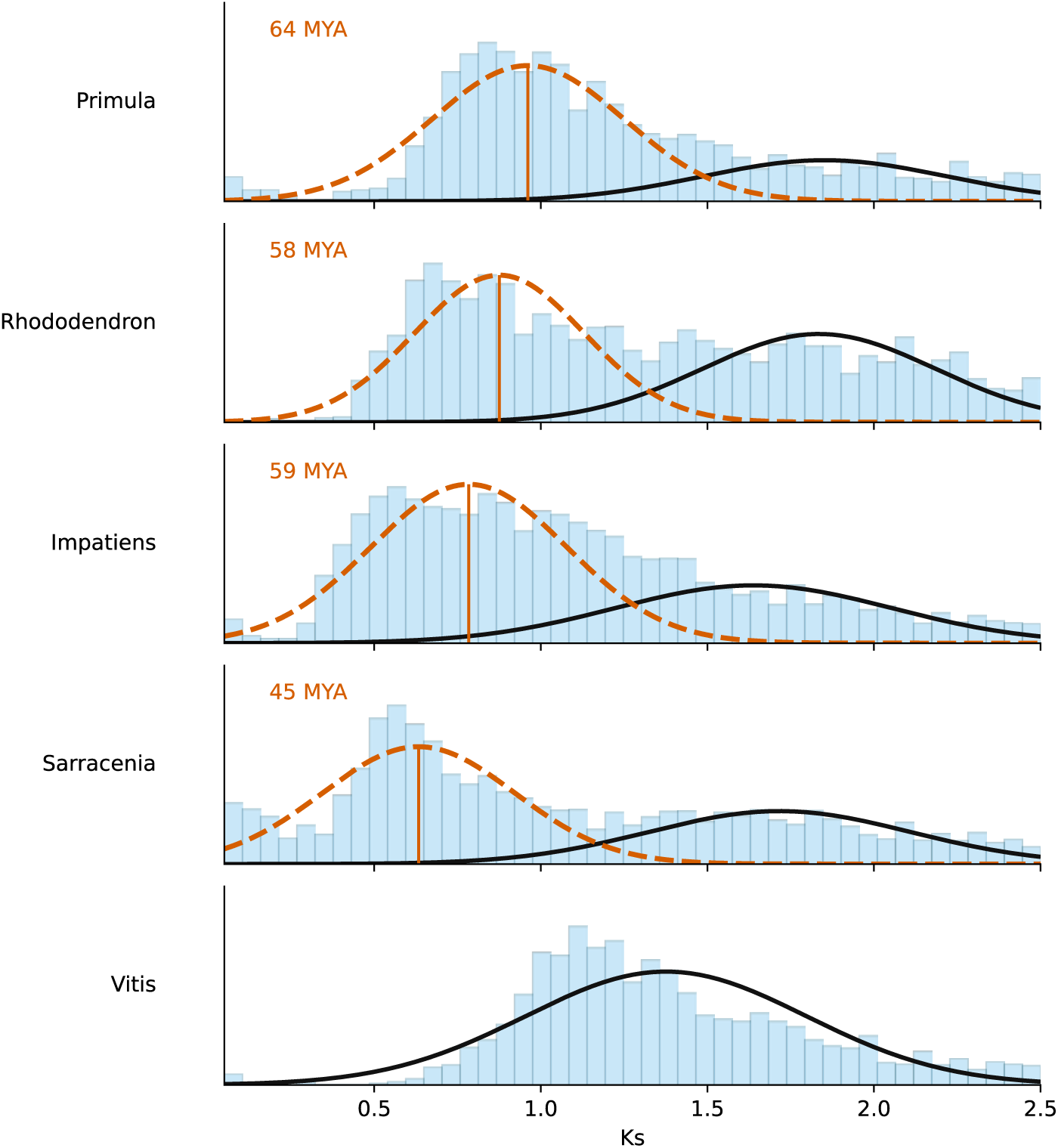
Synonymous substitution (Ks) distributions and inferred duplication components across selected Ericales. Histograms show Ks densities for self-syntenic duplicate gene pairs in *Sarracenia purpurea*, *Rhododendron*, *Impatiens*, *Primula*, and *Vitis vinifera* (0.05-2.5). Solid curves denote the broad, oldest Gaussian mixture component, interpreted as retention from the ancient core eudicot *gamma* hexaploidy, which is present in all taxa shown. Dashed curves indicate younger Ks components inferred from mixture modeling. *Vitis*, which has a comparatively slow molecular evolutionary rate, is shown as an external reference and exhibits a single retained component in this Ks range. Solid vertical lines mark the modal Ks values of the younger components, for which associated ages were calibrated independently within each species using their corresponding *gamma* component as a reference. Together, these profiles indicate widespread retention of deep duplications across Ericales, while leaving unresolved whether the younger Ks modes represent homologous or independent polyploidy events among lineages.

The proximity of these inferred ages raises the possibility of broadly contemporaneous duplication activity during the Paleocene to early Eocene. However, differences in Ks peak shape, breadth, and prominence, together with sensitivity to mixture modeling and assembly quality, preclude confident assignment of homology among these events. We therefore treat these post-*gamma* components as lineage-specific, or potentially shared, WGDs without asserting a single Ericales-wide polyploidy event.

### 2.2 Syntenic Architecture and Subgenome Structure Relative to *Vitis*

Comparative synteny across Ericales confirms that the *Sarracenia purpurea* genome retains a deeply conserved chromosomal backbone despite extensive lineage-specific rearrangement and fractionation. Anchoring against multiple Ericales genomes, including *Actinidia*, *Vaccinium*, *Rhododendron*, and *Impatiens*, reveals large blocks of conserved gene order that trace back to shared ancestral genome. These patterns establish *Sarracenia* as firmly embedded within the Ericales genomic framework rather than representing a structural outlier. To place these relationships within a common evolutionary reference, we next used *Vitis vinifera* as an outgroup to resolve gene-duplication ancestry and chromosomal correspondence. Syntenic mappings of *Sarracenia* chromosomes against *Vitis* as the target reference reveals, via fractionation bias differences^40,48^, long stretches of syntenic matching, largely at a 2:1 ratio (**Figure 8A**). In parallel, visualization of the same *Vitis*-referenced coordinate system across multiple Ericales taxa provides a structural overview of shared chromosomal correspondence that frames the more detailed analyses below (**Figure 8B**).

**Figure 8.**
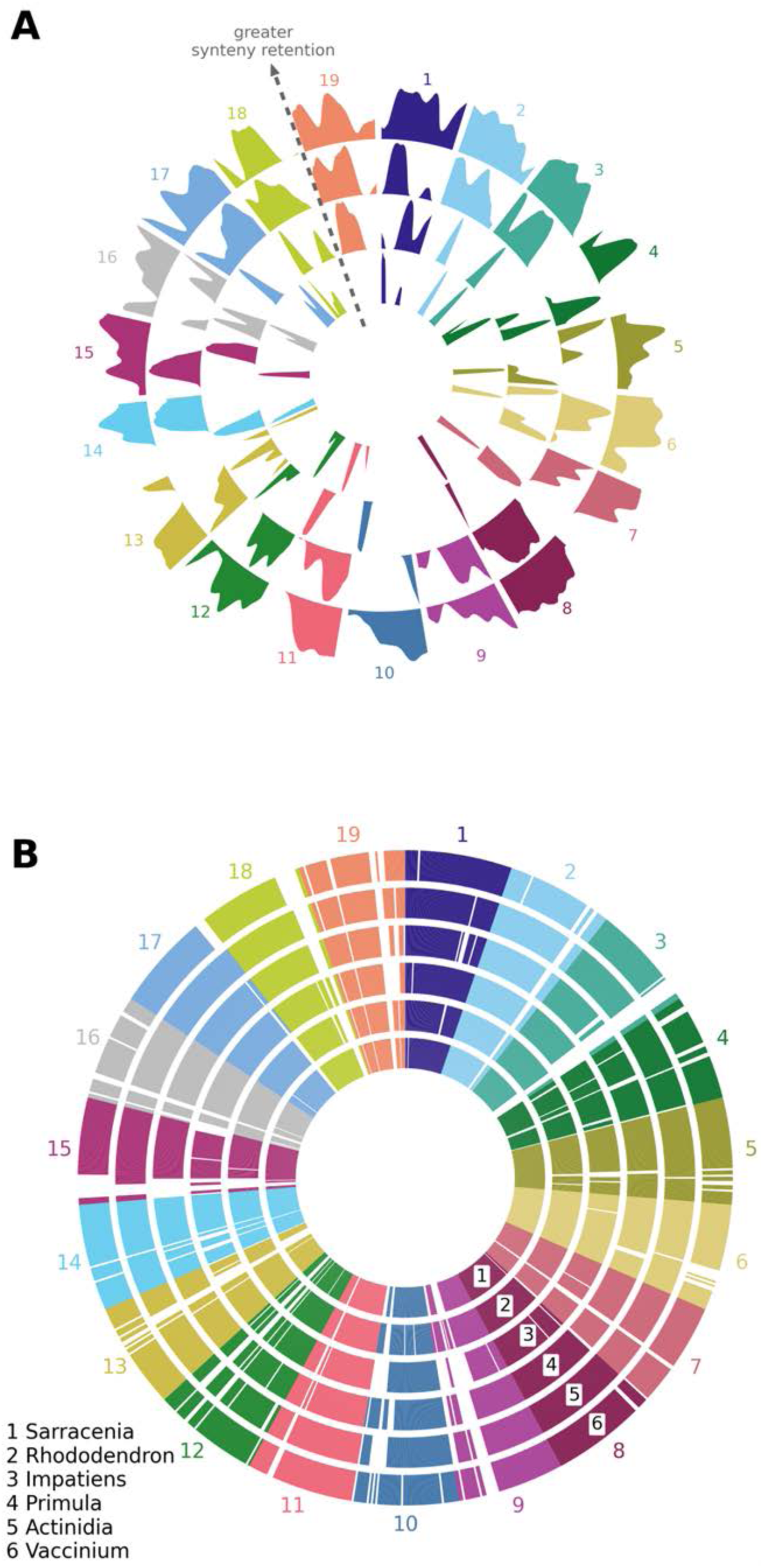
Radial representation of blockwise synteny retention and fractionation bias between *Sarracenia purpurea* and *Vitis vinifera*. Each colored sector corresponds to a single *Vitis* chromosome (1–19), arranged circularly. For each chromosome, the upper boundary of each colored band explicitly traces relative synteny retention (fractionation bias) along that chromosome, summarized across fixed bins, while the filled region beneath this boundary highlights the local magnitude of retention within the same syntenic layer for visual continuity. The dashed guide indicates increasing synteny retention within a given chromosome-layer representation, rather than comparisons among chromosomes or layers. The irregular, alternating profiles illustrate that regions of higher and lower retention are interleaved along chromosomes, consistent with a mosaic pattern of dominant and recessive fractionation rather than chromosome-wide uniformity. For each *Vitis* chromosome, up to four concentric syntenically anchored layers were recovered from the four best-matching *Sarracenia* segments; the outer filled ribbon reflects the strongest match and the inner ribbons progressively weaker matches. Nearly all *Vitis* chromosomes exhibit two strong outer matches from *Sarracenia*, consistent with the expected paleotetraploid history of the lineage. Additional inner ribbons identify weaker or more fragmented syntenic contributions and often correspond to structurally rearranged or fractionated regions inferred from subgenome analysis. A subset of the innermost segments may represent residual signal from the ancestral *gamma* hexaploidy shared broadly across core eudicots. (**B**) Representation of coordinated syntenic anchor presence and continuity across Ericales species mapped onto the same *Vitis* reference chromosomes. Concentric rings correspond to *Sarracenia*, *Rhododendron*, *Impatiens*, *Primula*, *Actinidia*, and *Vaccinium* (inner to outer), with each ring encoding per-bin syntenic anchor presence along *Vitis* chromosomes. This panel emphasizes the shared chromosomal coordinate framework across Ericales while illustrating lineage-specific variation in anchor presence and continuity, independent of fractionation-bias magnitude shown in panel **A**.

Furthermore, more detailed *Sarracenia-Vitis* synteny comparison shows alternating stretches of high and low syntenic anchor density along chromosomes. These intervals correspond to dominant and recessive subgenomic segments (**Figure 9**), as defined by fractional retention patterns (**Figure 8A**). Dominant segments preserve dense arrays of *Vitis*-derived anchors, whereas recessive segments show reduced anchor density and extended intervals with sparse syntenic signal (**Figures 8-10**). **Figure 10** quantifies these patterns explicitly as blockwise variation in anchor retention along reference chromosomes, without invoking explicit syntenic path reconstruction. Recessive segments also tend to occur in repeat-rich genomic contexts, consistent with a strong negative association between syntenic anchor density and transposable element abundance across 1 Mb genomic windows (Spearman’s ρ ≈ −0.5, P ≪ 0.001; see section 2.1.3).

**Figure 9.**
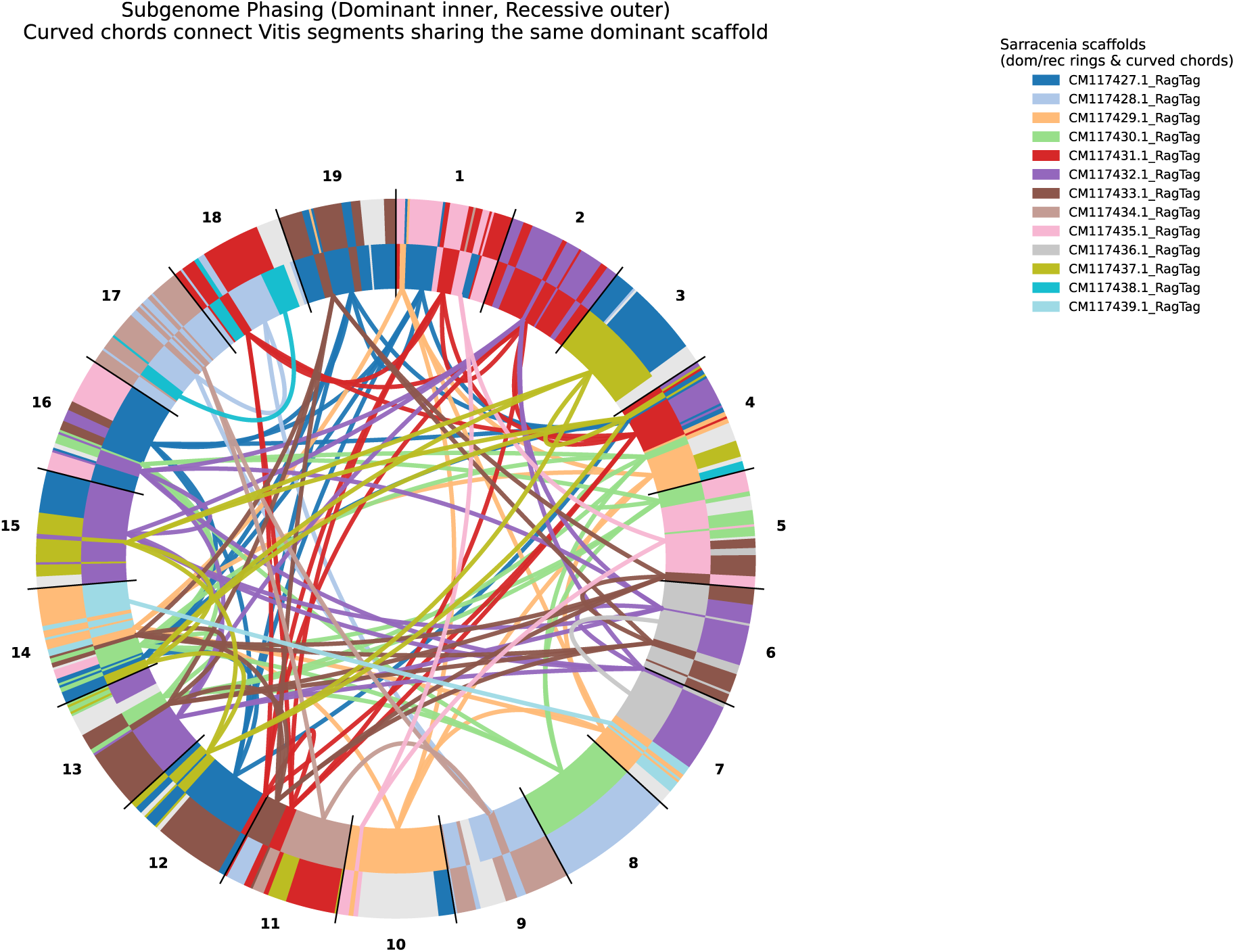
Predicted subgenome structure in the *Sarracenia purpurea* genome is accompanied by interleaved mosaicism against the *Vitis vinifera* genome. A circular representation of *Sarracenia* chromosomes syntenically mapped onto the *Vitis* genome (whose 19 chromosomes are numbered) reveals generally dominant (inner ring) and recessive subgenome (outer ring) predictions based on fractionation patterns. The subgenomes are assigned blocks colored by *Sarracenia* pseudochromosome, and the colored arcs connect dominant blocks (colored by respective pseudochromosomes), illustrating non-one-to-one correspondence and rearranged relationships among ancestral chromosomal segments relative to *Vitis*. Number of bins (*B*) used was 50 (see Methods section 4.3.2).

**Figure 10.**
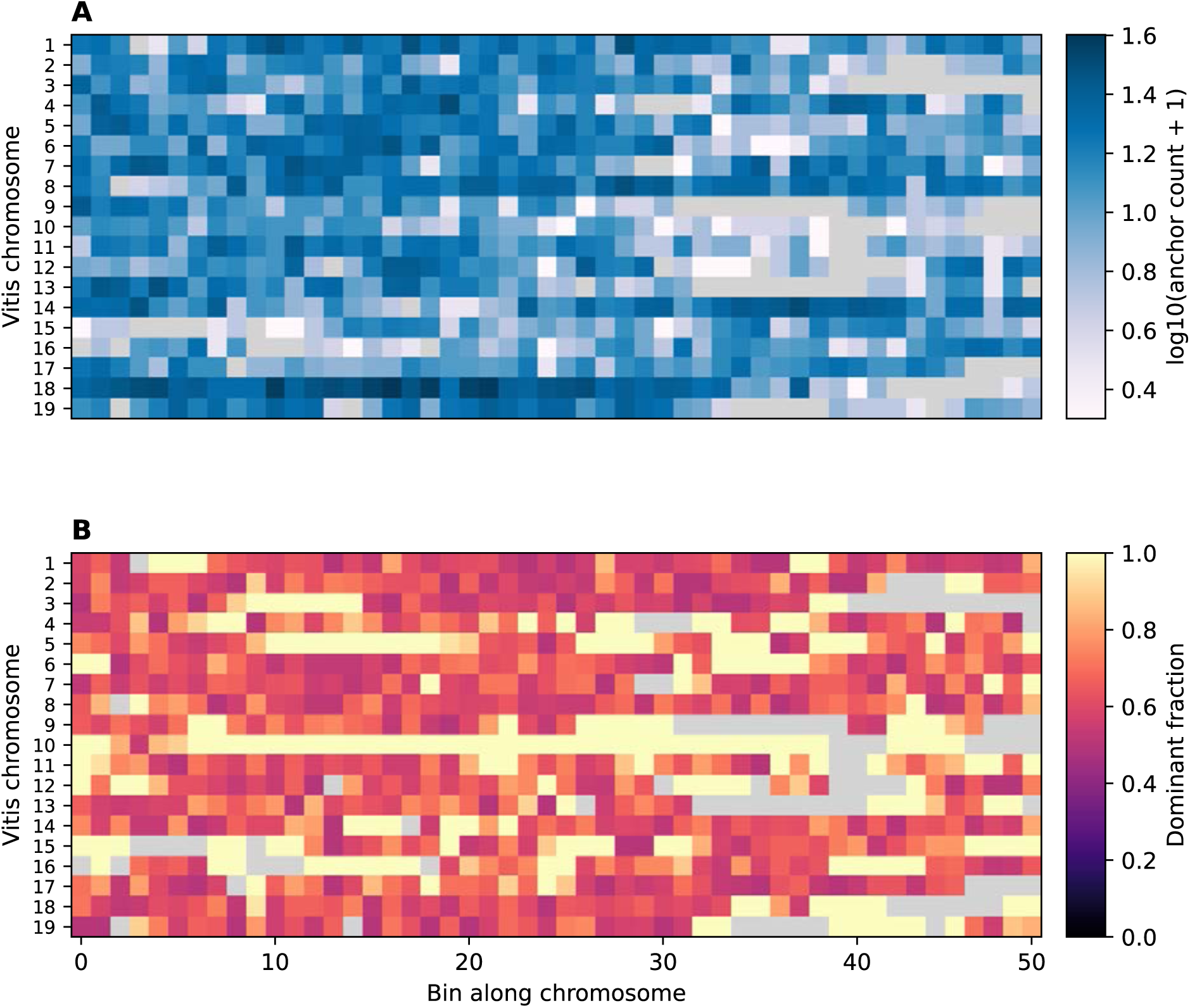
Blockwise retention heatmaps for *Sarracenia*-*Vitis* synteny. **(A)** Heatmap showing log₁₀-scaled syntenic anchor counts for 50 bins along each *Vitis* chromosome. A sequential blue-white-violet color scale indicates increasing syntenic anchor density. The pattern reveals alternating domains of high and low retention, with no chromosome-level uniformity, indicating that conserved and fractionated regions are tightly interwoven. **(B)** Heatmap depicting the fraction of dominant subgenome anchors per bin for the same *Vitis* chromosomes. Dominant-trending regions are marked by warm yellow hues, whereas more recessive regions transition through red to dark violet. The interspersed arrangement of color blocks demonstrates that subgenome segments tend to alternate between more dominant vs. recessive along chromosomes, forming a structural mosaic created by fractionation and localized rearrangements. Gray cells indicate the absence of syntenic anchors.

Blockwise retention heatmaps quantify these patterns explicitly. Across *Vitis* chromosomes, *Sarracenia* anchor density varies in a patchwork of high- and low-retention segments rather than showing chromosome-scale uniformity (**Figure 10A**). Regions of consistently high anchor signal form narrow bands corresponding to the most conserved subgenomic segments. Adjacent bins often transition abruptly to lower-retention compartments, revealing erosion or rearrangement of WGD-derived structures. These alternating intervals indicate that dominant and recessive segments are interleaved along chromosomes, rather than partitioned into intact, chromosome-wide domains (**Figure 10B**). The resulting mosaic pattern reflects differential fractionation and localized rearrangements accumulated over time, as captured by blockwise retention metrics rather than by explicit syntenic path inference.

### 2.3 Functional Architecture from Venn Bins in the Nine-Taxon Framework

To investigate how different classes of duplicated genes contribute to carnivorous traits in *Sarracenia*, a nine-taxon comparative framework was constructed to distinguish carnivorous innovation from background phylogenetic signal. Taxa incorporated were *Sarracenia*, *Nepenthes*, *Pinguicula*, *Utricularia*, *Actinidia*, *Vitis*, *Arabidopsis*, *Mimulus* and *Beta*. *Sarracenia’s* closest non-carnivorous Ericales relative in our sample^49^, *Actinidia*, provided a reference for identifying functional categories that belong to the Ericales heritage rather than to pitcher biology. *Mimulus* and *Beta* were included as non-carnivorous controls for the Lamiales (*Utricularia* and *Pinguicula*) and Caryophyllales (*Nepenthes*) carnivorous lineages, respectively^49^, ensuring that carnivore-associated signatures were not simply reflections of background family-level genomic tendencies. *Vitis* served as a deep structural outgroup, while *Arabidopsis* supplied standardized homolog identifiers and a high-quality GO annotation system.

Two distinct operations are used in this framework and must be kept conceptually separate. First, the nine-taxon bins are defined by strict intersections of *Arabidopsis*-mapped homolog presence and absence across taxa, so that each bin contains only those homologs present in exactly the taxa specified by its occupancy pattern and absent from all others. Second, GO enrichment is performed on functional foregrounds assembled as unions of bins for a given lineage, or as unions of enriched GO term sets across lineages, because convergence is expected to recur most reliably at the level of biological processes even when the underlying gene families differ.

We first annotated every *Sarracenia* tandem and syntenic duplicate in terms of its closest *Arabidopsis thaliana* homolog. The same mapping was generated for the other eight taxa: *Nepenthes*, *Pinguicula*, *Utricularia*, *Actinidia*, *Vitis*, *Arabidopsis*, *Mimulus*, and *Beta*. For each *Arabidopsis* locus appearing in at least one species, we recorded a nine-element presence-absence pattern specifying in which taxa that homolog was detected. This procedure produced a fixed universe of gene bins, where each bin corresponds to a precise nine-taxon occupancy pattern (for example, present in *Sarracenia* and *Nepenthes* only; present in four carnivores but absent elsewhere; present in all Lamiales members; etc.). All duplicate genes from *Sarracenia* could then be assigned unambiguously to these bins. Because bin membership is defined strictly by intersections, the bins serve as a stable phylogenetic scaffold on which functional comparisons are built.

The functional analysis required GO enrichment, and this step operates on collections of bins rather than on individual bins. For example, to evaluate *Sarracenia*-specific genomic novelty, we first identified all bins whose occupancy pattern included *Sarracenia* and excluded all other species; the genes contained in the union of these bins formed the foreground for GO enrichment. Thus, the genomic novelty step uses gene-set intersections to define bins, but unions of bins to define candidate functional foregrounds (**Figure 11**).

**Figure 11.**
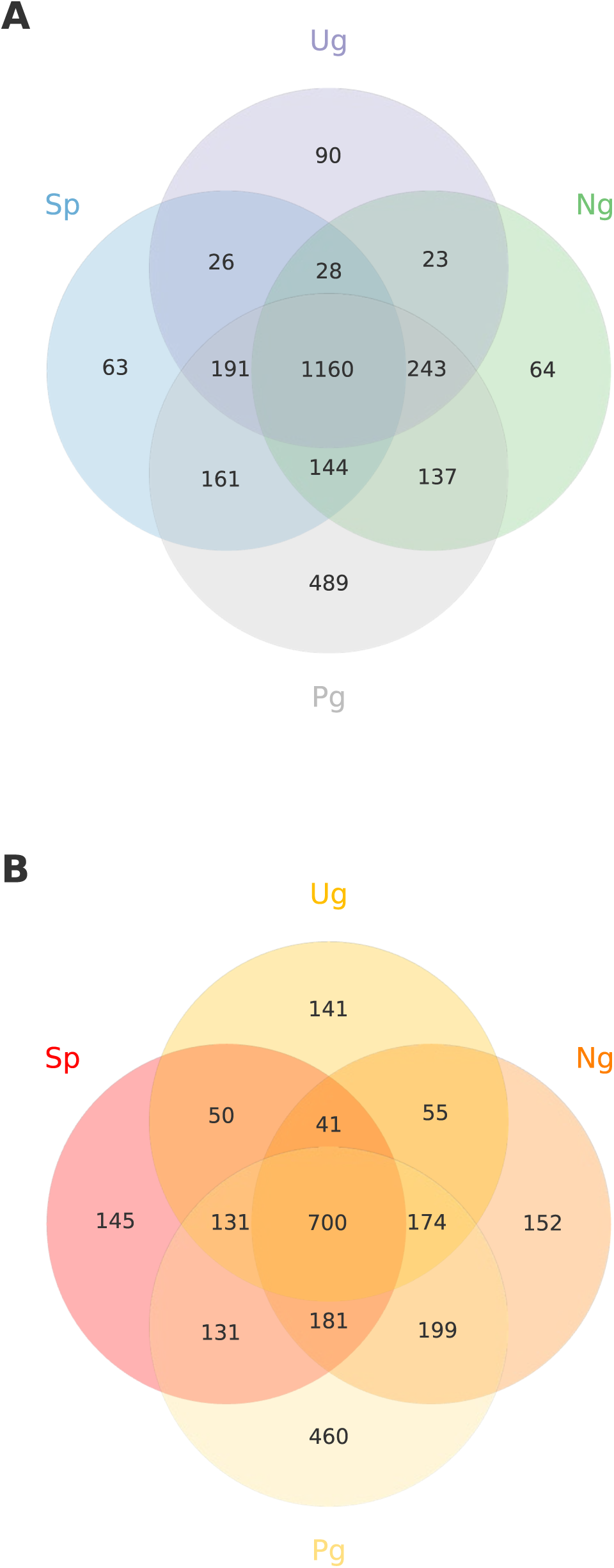
Overlap of enriched GO Biological Process categories across four carnivorous plant lineages. Venn diagrams summarize the number of significantly enriched GO Biological Process (BP) terms identified from duplicate-gene foregrounds in *Sarracenia purpurea* (Sp), *Nepenthes gracilis* (Ng), *Utricularia gibba* (Ug), and *Pinguicula vulgaris* (Pg). **(A**) GO BP terms enriched among syntenically retained duplicates, representing functions associated with ancient, dosage-constrained gene retention. **(B)** GO BP terms enriched among tandemly duplicated genes, representing functions associated with lineage-specific gene family expansion. Numbers indicate counts of distinct GO BP terms passing the same statistical threshold within each lineage and their intersections. Differences between panels highlight contrasting functional architectures associated with syntenic versus tandem duplication modes across independently evolved carnivorous plants.

The pitcher-associated analysis required integrating signals from two independently evolved pitcher-forming lineages, *Sarracenia* and *Nepenthes*. Because pitcher biology in these lineages likely involves distinct molecular strategies that may not converge at the level of individual gene families, we asked a functional question: which biological processes show significant enrichment in either lineage? Formally, the pitcher-associated functional space is the union of GO terms enriched in *Sarracenia* and in *Nepenthes*. The same logic was applied to reconstruct pan-carnivorous functional structure across *Sarracenia*, *Nepenthes*, *Pinguicula*, and *Utricularia*.

To distinguish carnivorous innovations from inherited Ericales background, *Actinidia* provided a necessary filter. Any GO category enriched in both a *Sarracenia*-containing foreground and in *Actinidia* was removed from the *Sarracenia* genomic novelty set, thereby isolating functional signals unlikely to reflect background Ericales biology.

Together, these procedures define a coherent comparative logic: intersection-defined bins provide the evolutionary structure, union-defined functional sets capture innovation and convergence, *Actinidia* filtration removes background Ericales signal, and pitcher-expression analyses identify which functional modules are actively deployed in trap tissues.

### 2.4 Comparative Functional Deployment Across Carnivorous and Ericales Lineages

#### 2.4.1 *Sarracenia*-Specific Functional Innovations

Comparative foreground analyses identify detoxification, redox, antifungal, and cuticle-modifying activities associated with the *Sarracenia* lineage, including both *Sarracenia*-only and carnivore-union categories (**Table S3**). These functions recur across duplication modes and include numerous tandem-derived loci. In addition to well-known effectors such as *PEN3*-like ABC transporters^50^, *GSTF*-related enzymes^51^, *CYP86*^52^- and *LACS2*^53^-associated cuticle-biosynthetic genes, the comparative foregrounds summarized in **Table S3** include other functionally interpretable homologs that carry defense-response GO terms representative of the functional categories captured in these bins. Collectively, these categories point to an expanded antifungal and xenobiotic-handling repertoire in *Sarracenia* relative to comparator lineages.

The *Sarracenia*-specific homolog of *DML1*^54^, which maintains stress-responsive gene accessibility by preventing inappropriate DNA methylation, appears among these functionally annotated, tandem-derived loci, highlighting chromatin-level regulatory processes within the *Sarracenia* innovation set summarized in **Table S3**.

These functional categories are consistently recovered in the *Sarracenia*-only innovation foregrounds and carnivore-union modules, although several overlap with *Actinidia* and are therefore excluded under the strict *Actinidia*-based filter. Six biological-process GO categories, response to oxidative stress, response to wounding, ethylene-activated signaling, cell differentiation, root-hair cell differentiation, and mitochondrial mRNA modification, are present in *Sarracenia* but absent from *Actinidia*, as summarized in **Table S3**. Detoxification, glutathione-based redox processes, antifungal functions, and cuticle-associated terms, although central to the broader *Sarracenia* innovation landscape, are shared with *Actinidia* and therefore are excluded under this filter.

In addition to these *Sarracenia*-only patterns, the comparative framework highlights uniquely enriched functional modules in the three other carnivorous lineages. Across *Nepenthes*, *Pinguicula*, and *Utricularia*, lineage-specific innovation sets emphasize secretory and vesicle-trafficking processes in *Nepenthes*, osmotic and desiccation-related pathways in *Pinguicula*, and rapid cell-wall remodeling and aquatic-habitat metabolic adjustments in *Utricularia*. These contrasts reinforce that, while the four carnivorous genera share a core stress-, interaction-, and metabolism-associated repertoire, each lineage exhibits distinct functional emphases captured by the comparative innovation framework. Relative to these patterns, *Sarracenia*’s unique signature is characterized by the oxidative, wound-responsive, and ethylene-associated biological processes summarized in **Table S3**.

#### 2.4.2 *Sarracenia*-*Nepenthes* Shared Pitcher Toolkit

The *Sarracenia-Nepenthes* pitcher comparison addresses how two independently evolved pitfall traps make use of duplicate-gene repertoires. Because this comparison tests convergence in enriched biological processes and does not require that the same *Arabidopsis*-mapped homologs fall in the strict Sp+Ng intersection bin, the relevant foreground comprises the union of GO terms enriched in either lineage. Under this union-based framework (**Table SB**), both taxa show significant enrichment for oxidative processes, cell-wall remodeling, and epidermal patterning functions, reflecting shared structural and ecological pressures associated with forming a tubular trap capable of accommodating fluid and degrading arthropod prey.

Although the two lineages differ in digestive strategy - *Nepenthes* secreting its own hydrolytic enzymes and *Sarracenia* relying more strongly on microbial activity - the overlap in enriched GO categories indicates convergent deployment of stress-responsive, cuticle-related, and cell-wall-modifying pathways. These shared pathways likely support the biomechanical and defensive requirements of constructing large, photosynthetically active, yet digestively functional pitcher leaves. Notably, convergence is observed at the level of functional categories rather than through a shared set of specific developmental regulators, consistent with the expectation that distantly related carnivorous lineages recruit distinct but functionally analogous genes to achieve similar morphological outcomes.

#### 2.4.3 Deep Carnivorous Signatures Across Four Independent Lineages

A suite of oxidative and biotic-interaction processes emerges from the union of GO categories enriched across *Sarracenia*, *Nepenthes*, *Pinguicula*, and *Utricularia* (**Table SC**). These pan-carnivore foregrounds incorporate numerous *Arabidopsis* homologs (e.g., AT1G15520, AT2G29430, AT4G39840) whose annotations span defense response, interspecies interaction, and microbial-stimulus management. The recurrence of these processes across four phylogenetically distant carnivorous lineages suggests that leaf-derived traps repeatedly draw upon stress-response and microbe-associated molecular architectures as foundational components of prey capture and digestion.

Within this shared functional framework, individual lineages nevertheless emphasize different biochemical strategies. *Sarracenia* shows disproportionate enrichment of detoxification and redox-buffering modules, *Nepenthes* more strongly represents secretory and vesicle-trafficking processes, and Lamialean carnivores additionally recruit pathways associated with osmotic regulation and cell-wall adjustment (**Tables SD-SF**). Despite these differences, the overlapping foregrounds point to a common reliance on stress-responsive, interaction-mediating, and cell-wall-modifying capacities. Collectively, these functions delineate a molecular repertoire that plants repeatedly mobilize during the evolutionary transformation of leaves into specialized organs for prey acquisition and nutrient absorption.

#### 2.4.4 Distinguishing *Sarracenia* Innovations from Ericales Background

Comparative filtration using *Actinidia* narrows the *Sarracenia-specific* genomic signatures by excluding GO terms that reflect ancestral Ericales biology. Because *Actinidia* is a noncarnivorous Ericales relative lacking pitcher morphology and digestive physiology, GO categories shared between *Sarracenia* and *Actinidia* most likely represent background Ericales functions rather than carnivory-related innovation.

Six biological-process GO categories: response to oxidative stress, response to wounding, ethylene-activated signaling, cell differentiation, root-hair cell differentiation, and mitochondrial mRNA modification, are uniquely enriched in *Sarracenia* and absent from the *Actinidia* enrichment profile (**Table SG**). These terms define a regulatory and stress-integration axis that has been elaborated along the *Sarracenia* lineage and may underlie aspects of pitcher development and physiology.

By contrast, detoxification, glutathione-dependent redox activity, antifungal responses, cuticle modification, and broader transport-related functions - although central to pitcher biology - do not survive the *Actinidia* filter. Their removal indicates that these pathways reflect ancestral Ericales capabilities rather than uniquely derived innovations.

Together, these filtered signatures emphasize that *Sarracenia* carnivory arose through targeted redeployment and elaboration of regulatory and stress-integration functions, rather than through wholesale invention of new biological processes. Only a narrow set of enriched regulatory categories remains uniquely associated with the *Sarracenia* lineage after Ericales background is removed (**Table S3**).

#### 2.4.5 Expression Patterns Reinforce the Venn-Bin Structure

*Sarracenia* gene expression analyses follow the same conceptual goal as the genomic comparisons but necessarily use different inputs. Differentially expressed genes (DEGs)were identified between flowers and pitcher leaves, mapped to *Arabidopsis* homologs using the same orthology framework as the genomic analyses, and analyzed for GO enrichment. Because differential expression reflects regulatory deployment rather than phylogenetic occupancy, presence-absence bins were not applied; instead, enriched GO terms from the DEG sets were compared qualitatively with Venn-derived functional modules.

Transcriptome comparisons of flowers versus pitcher leaves show that many genes supporting the nine-taxon Venn structure are strongly expressed in traps. Within the full DEG set, multiple *Sarracenia* homologs of *PEN3*, *ZIP6*^55^, and *DML1* show significant trap-biased expression (negative FvL log₂FC), reflecting xenobiotic export, micronutrient handling, and redox-responsive chromatin modulation (**Tables SH and SI**) - processes fundamental to the digestive and microbially mediated physiology of pitcher leaves. Several unnamed but functionally coherent pan-carnivore homologs (e.g., AT1G01580, AT1G24140, AT2G16390) also show trap-biased expression for at least one paralog, reinforcing their inferred role in carnivorous plant innovation (**Tables SH, SI, and SC**). The heatmap and volcano plots (**Figs. 12 and 13**, respectively) highlight the most extreme FvL differences, whereas genes such as *PEN3*, *ZIP6*, and *DML1* are supported by consistent trap-biased expression across the broader DEG set.

**Figure 12.**
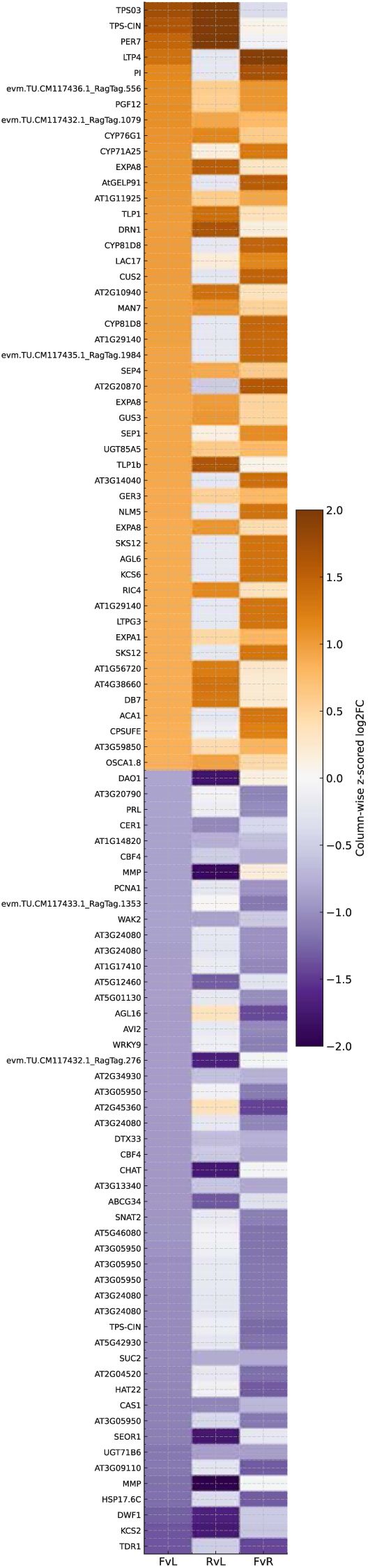
Differential expression heatmap for flowers versus pitcher leaves in *Sarracenia purpurea*. Heatmap showing log₂ fold-change values (flowers vs. pitcher leaves: FvL) for the top differentially expressed genes between flowers and pitcher leaves. Positive log₂ fold-change values (flower-biased expression) are shown in warm ochre tones, whereas negative values (pitcher-biased expression) are shown in cool blue tones, using a diverging color palette centered at zero. Genes are ordered by their FvL values, with flower-biased genes at the top and trap-biased genes at the bottom. For each *Sarracenia* gene, the best-supported *Arabidopsis* ortholog is shown - either by gene symbol when available or by *Arabidopsis* gene model ID when no symbol exists; *Sarracenia* gene IDs are shown only when no *Arabidopsis* homolog could be assigned (**Table SI**). The heatmap highlights canonical floral regulators such as *PI*, *SEP1*, and *SEP4*, along with terpene synthases (*TPS03*, *TPS-CIN*), peroxidases (*PER7*), and lipid-transfer proteins (*LTP4*) among the most strongly flower-upregulated genes. In contrast, pitcher-enriched genes include stress-response factors, cell-wall-modifying enzymes, hormone-associated regulators, and several uncharacterized but carnivore-associated homologs (**Tables SH and SC**), reflecting metabolic, structural, and physiological specializations of the trap.

**Figure 13.**
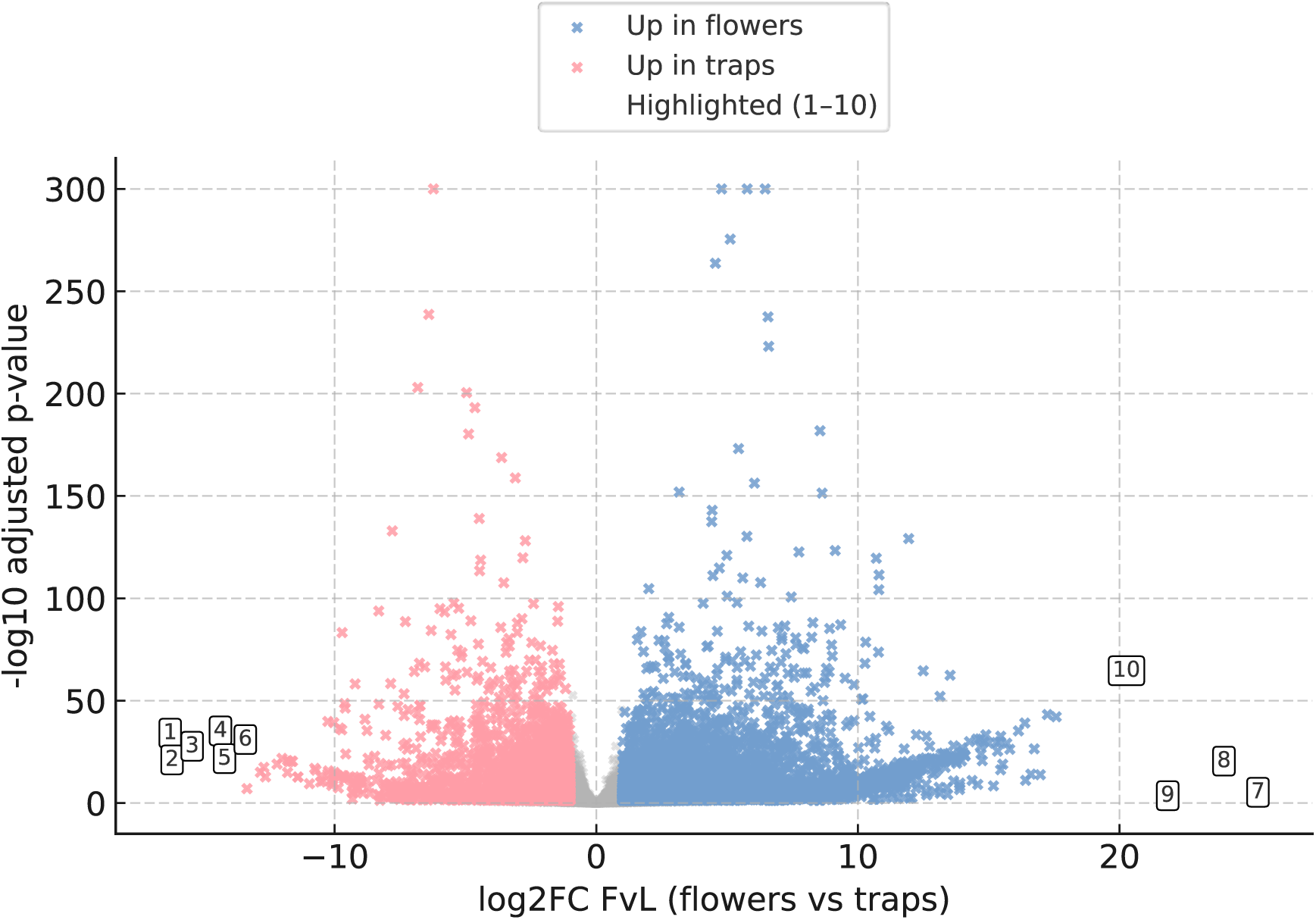
Differential expression volcano plot for flowers vs. pitcher leaves. Volcano plot showing differential gene expression between flowers and pitcher leaves in *Sarracenia purpurea*. For the expression contrast, positive log₂ fold-change values indicate higher expression in flowers, and negative values indicate higher expression in pitcher leaves (**Tables SA and SG**). Blue points denote significantly flower-upregulated genes (FDR < 0.05, |log₂FC| > 1), pink points denote trap-upregulated genes, and grey points are nonsignificant. Ten transcripts with the largest absolute expression shifts and informative functional annotations are highlighted and numbered (**1-10**, **Table SI**), with expression support summarized across innovation-related gene families (**Table SH**). **1:** *evm.TU.CM117430.1_RagTag.1674* / AT3G23230 (*TDR1*; Transcriptional Regulator of Defense Response 1) - encodes a member of the ERF (ethylene response factor) subfamily B-3 of ERF/AP2 transcription factor family, strongly trap-upregulated; **2:** *evm.TU.CM117436.1_RagTag.1249* / AT1G04220 (*KCS2*) - trap-biased 3-ketoacyl-CoA synthase, consistent with altered cuticular and lipid metabolism in pitchers; **3:** *evm.TU.CM117427.1_RagTag.1655* / AT3G19820 (*DWF1*) - trap-up brassinosteroid biosynthesis component, potentially modulating trap growth; **4:** *evm.TU.CM117432.1_RagTag.1718* / AT1G53540 (*HSP17.6C*) - trap-enriched small heat-shock protein, possibly indicative of stress-tolerant pitcher physiology; **5:** *evm.TU.CM117437.1_RagTag.1358* / AT1G70170 (*MMP*) - trap-up metalloprotease homolog, whose expression is nduced by fungal and bacterial pathogena; a plausible contributor to digestive or remodeling proteolysis; **6:** *evm.TU.CM117433.1_RagTag.1486* / AT3G09110 - strongly trap-biased gene of unknown function; **7:** *evm.TU.CM117428.1_RagTag.2440* / AT5G59310 (*LTP4*) - flower-biased lipid-transfer protein; **8:** *evm.TU.CM117427.1_RagTag.7* / AT1G30870 (*PER7*) - flower-up peroxidase; **9:** *evm.TU.CM117431.1_RagTag.1037* / AT3G25830 (*TPS-CIN*) and **10:** *evm.TU.CM117430.1_RagTag.70* / AT4G16740 (*TPS03*) - highly flower-enriched terpene synthases potentially contributing to floral volatile blends.

Flower-biased expression corresponds to terpene synthases (*TPS03*^56^, *TPS-CIN*^57^), peroxidases (*PER7*^58^), lipid-transfer proteins (*LTP4*^59^), cytochrome P450s (*CYP76G1*^60^, *CYP71A25*^61^), and canonical floral organ identity regulators (*PI*, *SEP1*, *SEP4*, *AGL16*^62,63^) (**Tables SH and SI**). Their flower-enriched deployment suggests that *Sarracenia* pitchers do not co-opt major floral-identity networks (at least at late stages of trap development, from which our RNA samples were isolated).

Trap-upregulated genes highlight molecular programs underpinning pitcher specialization. Strongly trap-biased transcripts include *TDR1*^64^, *KCS2*^65^, *DWF1*^66^, *HSP17.6C*^67^, *MMP*^68^, and additional stress- and metabolism-associated loci (**Tables SH and SI**). These reflect pathways associated with cuticle and wax biosynthesis, brassinosteroid metabolism, stress mitigation, osmotic buffering, temperature tolerance, and proteolytic remodeling - traits central to pitcher performance. Additional regulators such as *CER1*^69^, *CBF4*^70^, *WRKY9*^71^, *PCNA1*^72^, *PRL*^73^, and *DAO1*^74^ reinforce that pitcher identity integrates surface modification, stress-response, hormonal, and redox-balance modules (**Tables SH and SI**).

Flower-enriched defenses, including tandem-only homologs such as AT1G02205, AT1G55260, and AT1G62045, suggest that reproductive tissues maintain broader defense capacities attenuated in traps (**Tables SH and SI**). Distinct paralogs of *PEN3* and *ZIP6* occur in both trap-up and flower-up sets, indicating differential subfunctionalization of duplicated transporters across tissues.

Overall, these transcriptional patterns validate and extend the Venn-bin framework: many lineage-specific and carnivore-level innovations identified from genomic analyses are actively deployed in traps, while other functions remain primarily associated with vegetative or floral contexts. Pitchers emerge from a modular redeployment of stress-response, developmental, metabolic, and structural pathways superimposed on the Ericalean genomic architecture.

### 2.5 Integrated Model for the Evolution of *Sarracenia* Pitcher Biology

The combined syntenic, duplication-mode, and functional-foreground analyses outline a two-tiered model for pitcher evolution in *Sarracenia purpurea*. At its foundation lies a conserved regulatory scaffold housed preferentially on dominant subgenomes. Key developmental regulators, including *AGO1*^75^ (*ARGONAUTE 1*), *BRX*^76^ (*BREVIS RADIX*), *GATA11*^77^ (*GATA TRANSCRIPTION FACTOR 11*), *ETC1*^78^ (*ENHANCER OF TRY AND CPC 1*), and *RCD1*^79^ (*RADICAL-INDUCED CELL DEATH 1*), are retained as single or low-copy loci within high-retention syntenic blocks (**Table SI**), consistent with dosage sensitivity and long-term preservation following ancient polyploidy. The known developmental roles of these genes in *Arabidopsis*, spanning leaf polarity, epidermal cell-state regulation, auxin-linked patterning, and stress integrated developmental control, support a gene-level model in which pitcher morphology is constructed by redeploying conserved leaf developmental regulators rather than by invoking lineage-specific invention of entirely new regulatory pathways.

Superimposed on this scaffold is a suite of tandem expansions concentrated in structurally dynamic, often TE-rich genomic regions. These tandem-derived families are enriched for detoxification functions, glutathione-based redox buffering, antifungal responses, and cuticle-modifying and wall-remodeling processes (**Tables SA and SG**). Their functional signatures align closely with the biochemical and microbial demands of trap operation, particularly the need to regulate fluid chemistry, manage reactive metabolites, suppress fungal growth, and modify surface and luminal structures during prey decomposition.

To determine whether contrasting duplication modes are associated with differences in regulatory deployment, we compared expression responsiveness of genes retained through syntenic versus tandem duplication using pitcher-associated transcriptomic data. Tandem-derived genes show significantly greater expression responsiveness than syntenically retained genes (cf. ^80^), as assessed by a two-sided Mann-Whitney U test comparing absolute log-transformed expression change across conditions (p = 6.94 × 10⁻¹⁸; Cliff’s δ = 0.179; n = 934 tandem, 4,578 syntenic; **Figure 14**). Expression responsiveness was quantified as the absolute value of the log₂ fold change (|log₂FC|) between pitcher and flower tissues, computed after addition of a fixed pseudocount to raw expression values to avoid undefined ratios. Tandem-derived genes exhibit both higher median responsiveness and broader variance, whereas syntenic genes display narrower expression distributions consistent with stronger regulatory constraint.

**Figure 14.**
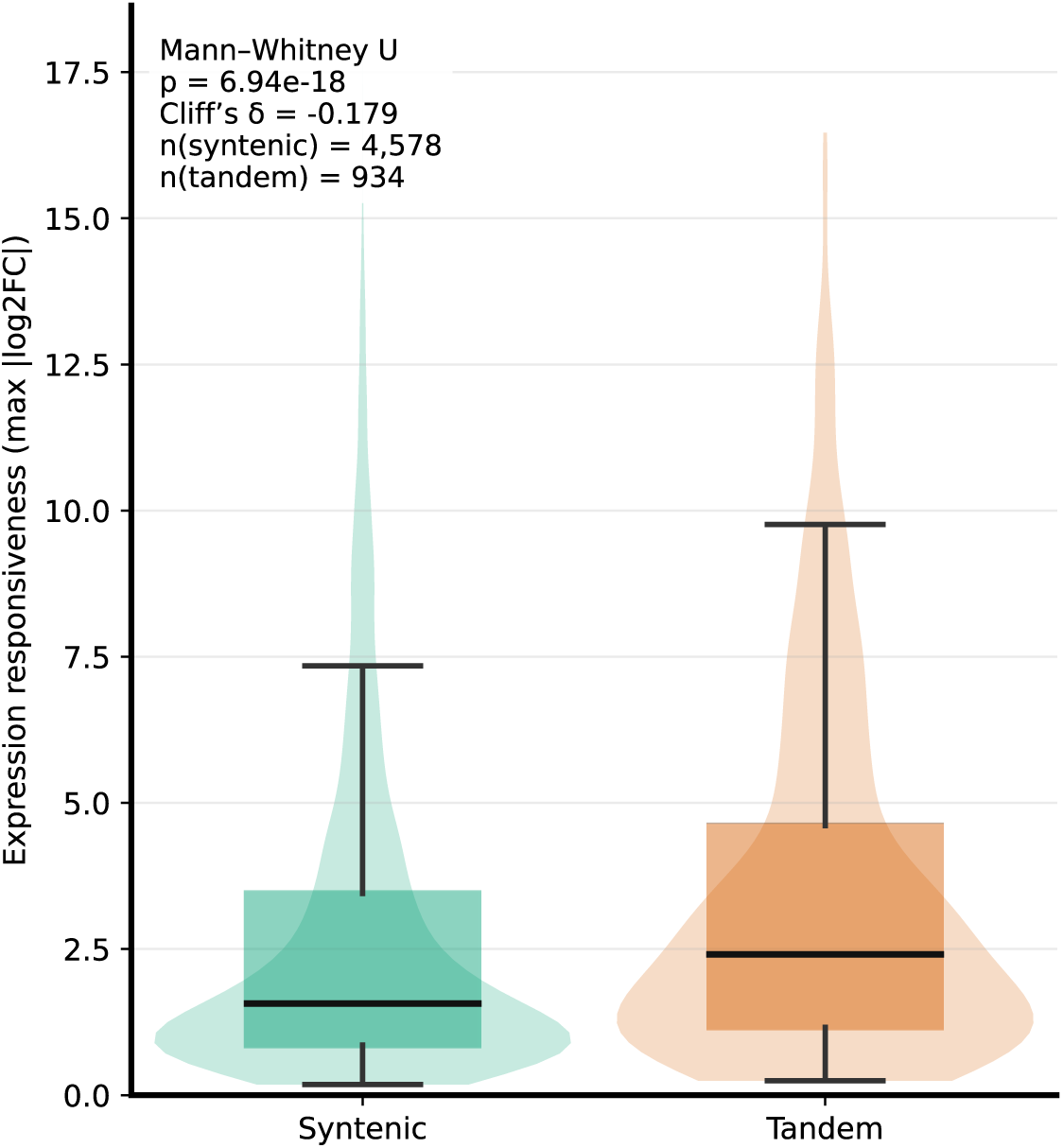
Expression responsiveness of syntenic and tandem-derived genes in *Sarracenia purpurea*. Violin and box plots summarize the distribution of expression responsiveness for genes retained through syntenic duplication versus tandem duplication. Expression responsiveness was quantified as the absolute log-transformed expression change across pitcher-associated transcriptomic conditions. Boxes indicate medians and interquartile ranges, while violins depict kernel density estimates of the full distributions. Statistical significance was assessed using a two-sided Mann-Whitney U test, with effect size reported as Cliff’s delta (δ). Sample sizes for each duplication class are indicated within the plot. Tandem-derived genes exhibit greater expression responsiveness and broader variance than syntenic genes, consistent with relaxed regulatory constraint in tandem-expanded gene families (**Tables SA and SH**).

We next asked whether these expression differences are accompanied by broader functional diversification. Functional breadth of syntenic and tandem gene sets was assessed using Gene Ontology (GO) biological-process annotations under a rarefaction framework that explicitly controls for unequal gene-set sizes. Across sampling depths, rarefaction curves for tandem-derived genes tend to exceed those for syntenic genes in both GO term richness and effective functional breadth (**Figure 15**). This pattern is consistent with tandem-expanded gene sets sampling a broader range of annotated biological processes when gene set size is controlled. Variability envelopes derived from repeated subsampling illustrate that this trend is consistent across iterations, although rarefaction is used here as a descriptive framework rather than for formal hypothesis testing. These results are consistent with tandem duplication contributing disproportionately to functional diversification in the *Sarracenia* genome relative to syntenically retained duplicates.

**Figure 15.**
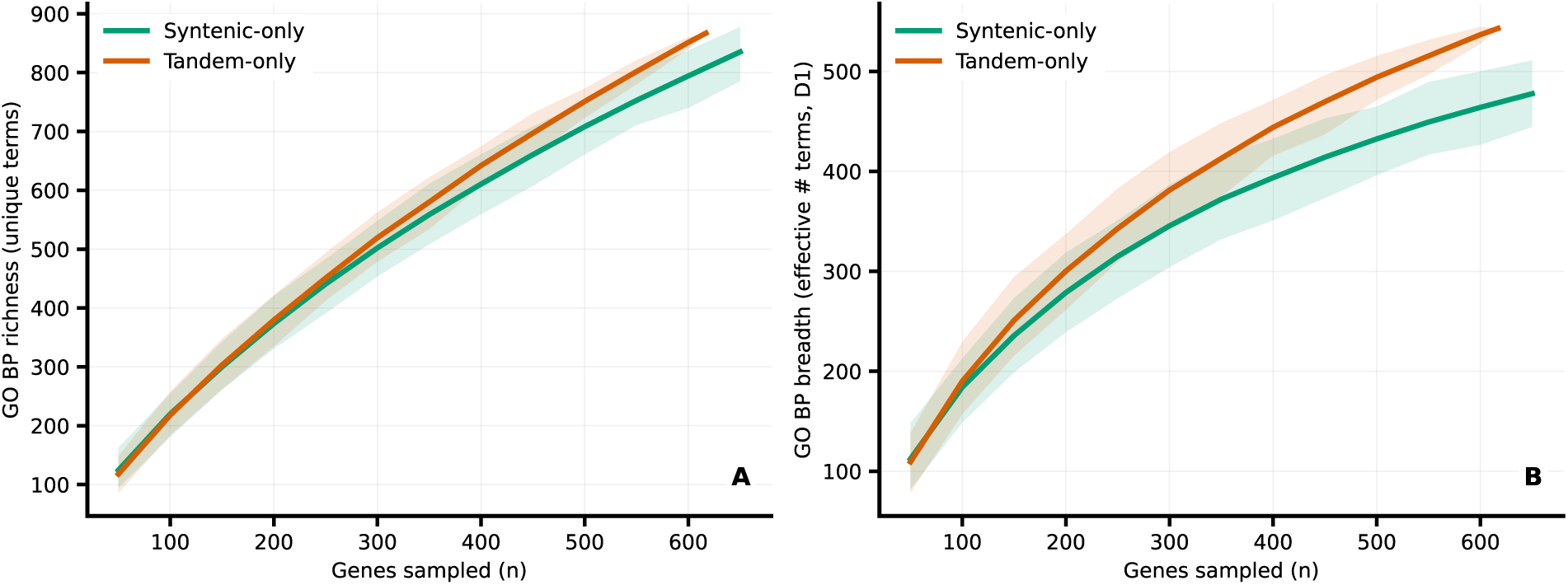
Functional breadth of syntenic and tandem gene sets inferred from GO rarefaction. Rarefaction curves depict functional breadth of syntenic and tandem gene sets based on GO biological-process annotations. Curves show mean GO richness and effective functional breadth (D1) as a function of subsampled gene number, with shaded envelopes indicating variability across subsampling iterations. Functional breadth was quantified using both the number of unique GO terms and the exponential of Shannon entropy (cf. ^81^). Rarefaction is used as a descriptive framework to control for differences in gene set size and visualize contrasts in functional breadth between duplication classes, without formal hypothesis testing across curves (**Tables SA and SC**).

Comparisons with *Nepenthes* reveal that, despite substantial differences in digestive strategy, the two lineages draw upon overlapping functional categories, including oxidative stress responses and wall-modifying pathways, that are enriched in both taxa under the union-based framework. This convergence occurs primarily at the level of functional deployment rather than strict gene-level sharing, reflecting independent recruitment of analogous molecular programs to achieve similar pitcher architectures.

Comparative timing of duplication and repeat landscapes in *Sarracenia* indicates that its lineage-specific WGD signal predates a much younger phase of LTR retrotransposon activity, placing large-scale gene duplication substantially earlier than the principal episode of repeat accumulation. Rather than acting as direct drivers of morphological novelty, LTR-rich regions can provide a structurally permissive genomic environment in which tandem duplication and local gene-family expansion can proceed, through mechanisms such as unequal crossing-over and local rearrangement, without disrupting dosage-sensitive regulatory networks.

Together, these patterns suggest that the evolution of *Sarracenia* pitchers involved (i) retention of a dosage-constrained developmental framework on dominant subgenomes, and (ii) diversification of effector modules through tandem expansion in more flexible genomic compartments. Subgenome asymmetry provides a mechanistic basis for this division of labor: dominant subgenomes preferentially retain dosage-sensitive regulators with little evidence of tandem proliferation, whereas recessive and boundary regions harbor many of the expanded detoxification, redox-related, and interaction-associated modules (**Tables SA, SG, and SI**). Because these latter functions may be more dosage flexible than WGD-surviving transcriptional networks, their expansion across genomic compartments could proceed without compromising developmental homeostasis, allowing the biochemical and microbial ecology of the trap to evolve rapidly while the core developmental program remained stable. This division of labor provides a mechanistic explanation for how regulatory stability and ecological specialization were simultaneously achieved during the origin of carnivorous leaf function.

## 3. Conclusions

The chromosome-scale genome assembly of *Sarracenia purpurea* provides an integrated view of the structural and functional processes that supported the origin of pitcher leaves by resolving how different modes of gene duplication contributed to distinct evolutionary roles. The genome retains extensive Ericales-derived syntenic structure and displays clear subgenome asymmetry, with dominant subgenomes enriched for dosage-sensitive developmental and signaling regulators that form a conserved architectural scaffold, and recessive compartments containing more flexible gene families. Tandem expansions in structurally dynamic regions contributed detoxification, redox, antifungal, and cuticle-associated activities that supply biochemical and ecological flexibility characteristic of the species’ microbially mediated digestive system.

Expression data reinforce this framework by linking genomic partitioning to regulatory deployment in pitcher tissue. Pitchers upregulate not only homologs of named *Arabidopsis* genes such as *PEN3*, *ZIP6*, and *DML1*, but also a broader suite of stress-associated, metal-handling, and wall-modifying loci. These additional genes match enriched *Sarracenia*-specific and carnivore-union GO categories and support a regulatory program integrating xenobiotic transport, micronutrient balancing, reactive-metabolite buffering, and targeted modification of epidermal and luminal surfaces. The presence of distinct paralogs in different Venn bins points to divergent regulatory trajectories among duplicates, suggesting that stress-integration modules were subdivided or refined as *Sarracenia* adapted to a semi-open, microbially rich digestive environment.

Comparative analyses across carnivorous taxa reveal that pitcher evolution repeatedly draws upon shared developmental and stress-responsive capacities, while functional diversification tracks the ecological demands of each trap type. Parallel enrichment of stress-, interaction-, and wall-modifying pathways across carnivorous lineages underscores a convergent functional basis for trap formation, whereas differences in effector composition highlight lineage-specific strategies for prey breakdown and nutrient acquisition.

*Actinidia*-based filtration indicates that only six GO biological-process categories - response to oxidative stress, response to wounding, ethylene-activated signaling, cell differentiation, root-hair cell differentiation, and mitochondrial mRNA modification - are uniquely enriched in *Sarracenia* relative to (at least some) other Ericales. In contrast, detoxification, glutathione-based redox buffering, antifungal functions, cuticle-associated pathways, and transport-related activities likely reflect lineage-specific elaboration of ancestral Ericales capacities rather than de novo innovation, underscoring how carnivory arises through redeployment and expansion of pre-existing genomic programs.

Together, these results demonstrate how ancient polyploidy vs. tandem duplication can partition regulatory and effector functions in ways that facilitate the evolution of novel organs. Conserved regulators on dominant subgenomes provide morphological and developmental stability, while tandem expansions supply the metabolic and ecological specialization required for a microbially mediated digestive environment.

A future chromosome-scale genome for *Cephalotus* will enable direct testing of how many components of the shared pitcher toolkit represent true molecular convergence rather than distinct elaborations of different genomic substrates. The *Sarracenia* genome thus anchors a broader understanding of carnivorous plant evolution by integrating structural legacy, regulatory conservation, and effector diversification into a unified model of trait innovation.

## 4. Methods

### 4.1 Genome Resources and Analytical Scope

All analyses were conducted on a chromosome-scale genome assembly of *Sarracenia purpurea* comprising 13 pseudochromosomes (publicly available at https://genomevolution.org/coge/GenomeInfo.pl?gid=69349). All genomic coordinates, gene models, and syntenic relationships refer to this assembly and its associated primary protein-coding gene annotation (also freely available at the above CoGe link). Each gene locus was represented by a single representative protein-coding model. Genome-wide transposable element annotations were available as genomic intervals and were used only as positional features in downstream analyses. Gene density was quantified by counting gene midpoints in non-overlapping 1-Mb windows across pseudochromosomes larger than 5 Mb, using coordinates from the final gene annotation GFF.

### 4.2 Genome Construction and Annotation Workflow

#### 4.2.1 Plant Material, DNA Extraction, and Oxford Nanopore Sequencing

Plant material and voucher specimens for sequencing were collected from mature individuals of *Sarracenia purpurea* at Mendon Ponds Park, Rochester, New York (**Figure 1**). Species identification was performed in the field. Voucher specimens are currently deposited at the University at Buffalo. Leaf tissue was placed into individual vials and stored at −80 °C until DNA extraction.

Approximately 10 g of flash-frozen tissue was used for isolation of high-molecular-weight (HMW) genomic DNA. Nuclei isolation followed the BioNano NIBuffer protocol, in which frozen leaf tissue was homogenized in liquid nitrogen and subjected to a nuclei lysis step using IBTB buffer supplemented with spermine and spermidine and filtered immediately prior to use. IBTB buffer consisted of Isolation Buffer (IB; 15 mM Tris, 10 mM EDTA, 130 mM KCl, 20 mM NaCl, 8% (m/V) PVP-10, pH 9.4) containing 0.1% Triton X-100 and 7.5% (v/v) β-mercaptoethanol (BME), mixed and chilled on ice. The homogenized tissue-buffer mixture was strained to remove undissolved plant material. Triton X-100 was then added to a final concentration of 1% to lyse nuclei, followed by centrifugation at 2000 × g for 10 min to pellet the nuclei. Following nuclei isolation, DNA was extracted using a cetyltrimethylammonium bromide (CTAB) protocol modified for Oxford Nanopore sequencing. The quality and concentration of HMW genomic DNA were assessed using a NanoDrop spectrophotometer and agarose gel electrophoresis following standard protocols. Extracted genomic DNA was further purified using a Qiagen Genomic-Tip 500/G according to the manufacturer’s instructions. Aliquots of purified DNA were subjected to treatment with Short Read Eliminator (SRE; PacBio) to perform size selection and enrich for DNA fragments longer than 10 kb

High-molecular-weight genomic DNA was thereafter sequenced using Oxford Nanopore Technology (ONT) platforms. Library preparation and sequencing followed standard ONT protocols appropriate for high-molecular-weight plant genomic DNA. Sequencing generated long-read data without pre-filtering for read length or read quality. All genome assemblies and downstream analyses were performed using unfiltered ONT reads, unless explicitly stated otherwise, in order to preserve information from repeat-rich and structurally complex regions of the genome.

#### 4.2.2 Genome Assembly

Unfiltered Oxford Nanopore reads were assembled using hifiasm v0.25.0 with the --ont flag enabled, which activates the assembler’s long-read mode optimized for nanopore data. Assembly was performed to generate a collapsed primary (haploid) assembly, rather than phased haplotypes, reflecting the study’s focus on genome structure, gene content, and duplication history rather than allelic variation.

Assembly completeness was evaluated using BUSCO v5.4.3 with the *eudicots_odb10* dataset, which assesses the presence and copy number of conserved single-copy orthologs expected in eudicot genomes. Assembly contiguity and size statistics were evaluated using QUAST v5.2.0.

To assess read support and facilitate downstream curation, unfiltered ONT reads were aligned back to the initial assembly using minimap2 v2.24, employing presets appropriate for long-read alignment. Resulting alignments were sorted using SAMtools v1.16.1.

#### 4.2.3 Contamination Screening and Assembly Curation

The initial genome assembly was screened for potential contamination using BlobTools v1.1.1 in conjunction with BLAST+ v2.12.0. For each contig, taxonomic assignments were inferred from BLASTN similarity searches. Contigs classified as Streptophyta were retained, as were contigs lacking confident taxonomic assignment (no-hit or undefined). Contigs assigned to non-plant taxa were removed from further analyses.

Unfiltered ONT reads aligning to the retained contigs were extracted and reassembled using hifiasm with the same parameters as the initial assembly. This produced a curated assembly in which non-plant sequences were excluded while preserving read depth and long-range genomic structure across retained regions. Assembly quality and completeness were reassessed using BUSCO and QUAST as described above.

#### 4.2.4 Homology-Based Scaffolding

The curated assembly was scaffolded using RagTag v2.1.0, employing a publicly available *Sarracenia purpurea* genome assembly (NCBI accession GCA_051027295.1) as a reference guide for contig ordering and orientation. RagTag was used strictly in reference-guided scaffolding mode; nucleotide sequences were not modified, and no gap filling or sequence correction was performed.

This scaffolding step produced a chromosome-scale assembly comprising 13 pseudochromosomes and 1,871 unplaced scaffolds. All subsequent analyses were conducted using this RagTag-scaffolded genome assembly.

#### 4.2.5 Genome Annotation

Repetitive elements were identified using a combination of RepeatModeler v2.0.4 and EDTA v2.0.1, capturing both de novo repeat families and structurally defined transposable element classes. Repeat libraries generated by the two approaches were merged and collapsed into a non-redundant repeat library using CD-HIT v4.8.1, applying thresholds of 95% sequence identity and 90% alignment coverage.

Protein-coding gene prediction was performed using GeMoMa v1.9.0 with default parameters. Reference proteomes from five angiosperm species were used to guide prediction: *Arabidopsis thaliana*, *Camptotheca acuminata*, *Actinidia eriantha*, *Camellia lanceoleosa*, and *Vaccinium darrowii*. These species were selected to provide phylogenetic breadth within eudicots while remaining relevant to Ericales. RNA-seq data were incorporated as evidence for the prediction of splice sites.

This annotation was refined using EVidenceModeler (EVM) v2.1.0 to standardize naming and retain the best-supported primary gene model at each locus. The resulting EVM-integrated GFF3 annotation served as the final gene annotation used for all downstream analyses. Annotation completeness was assessed using BUSCO v5.5.0 on the predicted protein set.

### 4.3 Syntenic Anchor Processing and Coordinate Binning

#### 4.3.1 Syntenic Anchor Preprocessing

Syntenic anchors were provided as gene-gene correspondences between *Sarracenia purpurea* and reference genomes. These runs were executed in SynMap^39–41^ with FractBias^48^ enabled to generate standardized syntenic anchor sets against *Vitis* as a joint reference. While FractBias summary outputs were inspected for consistency, all fractionation metrics, dominance assignments, and downstream visualizations reported here were recomputed independently from the raw syntenic anchor tables. SynMap settings for FractBias were:

- DAGChainer: relative gene order = nucleotide distance; maximum distance between two matches = 20 genes; minimum number of aligned pairs = 5 genes
- Skip random/unknown chromosomes
- Syntenic depth: Quota Align^82^
- Ratio of coverage depth: *Vitis* = 1 against all other taxa = 4
- Overlap distance: 40
- Window size: 100 genes; use all genes in target genome

Each anchor record obtained from SynMap (at the link: “Results with synonymous/non-synonymous rate values”) contained identifiers for the source and target chromosomes, genomic coordinates for both genes, and a Ks value. FractBias results were downloaded at the “Fractionation Bias synteny report” and “Fractionation Bias sliding window results” links.

Anchors were filtered as follows:

- Anchors lacking defined chromosomal coordinates were excluded.
- Anchors mapping to unplaced or unanchored scaffolds were excluded.
- Anchors lacking Ks values were excluded from Ks-based analyses but retained for purely structural analyses, they were used for reference-coordinate projection across multiple Ericales taxa, where anchors were treated strictly as structural signals and were not used for dominance or fractionation-based inference.

#### 4.3.2 Reference-Coordinate Binning of Syntenic Anchors

All reference-based analyses were conducted in a fixed bin coordinate system.

This reference-coordinate framework was applied uniformly wherever syntenic anchor positions were summarized along the *Vitis* genome, including both lineage-specific and multispecies analyses, where anchors were interpreted descriptively as indicators of syntenic presence rather than quantitative density.

For each reference chromosome 𝑐:

Let 𝐿_𝑐_ be the maximum coordinate observed among all anchors on chromosome 𝑐.

Define a fixed number of bins 𝐵, held constant across all chromosomes for a given analysis. Define the bin width:

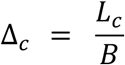

For each anchor with midpoint coordinate 𝑚 on chromosome 𝑐, assign the anchor to bin:

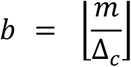

Clip 𝑏 such that:

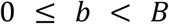

This produces a discrete bin index for every anchor, enabling direct comparison across chromosomes and taxa.

#### 4.3.3 Anchor Density Aggregation and Transformation

For each *Sarracenia* chromosome 𝑖 and reference bin 𝑗, the raw anchor count is:

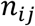

To stabilize variance and reduce domination by anchor-rich bins, counts were transformed as:

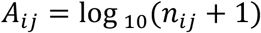

All subsequent analyses, including dominance, fractionation, path inference, and similarity matrices, operate exclusively on 𝐴*_ij_*. In analyses where anchors were summarized across taxa rather than partitioned by subgenome state, transformed per-bin anchor signals were interpreted descriptively as measures of anchor presence and continuity along the reference coordinate system.

### 4.4 Syntenic Retention Metrics and Summary Statistics

#### 4.4.1 Dominance Margin and Fractionation Bias

For each reference bin 𝑗, transformed anchor signals across chromosomes were ranked.

Let:

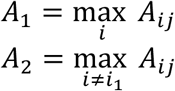

Where:

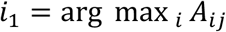

then the dominance margin is:

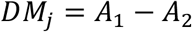

Bins were classified as:

- dominant if 𝐷𝑀*_j_* > 0
- recessive if 𝐷𝑀*_j_* ≤ 0

No magnitude cutoff beyond sign was applied.

Fractionation bias (see also^48^) was computed as:

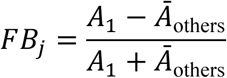

where *Ā*_others_ is the mean of all non-dominant transformed signals. Given 𝐴*_ij_* ≥ 0, then 0 ≤ 𝐹𝐵*_j_* < 1 whenever 𝐴_1_ ≥ Ā_others_.

#### 4.4.2 Syntenic Retention Similarity Matrices

Each *Sarracenia* chromosome was represented as a vector:

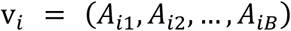

Pairwise similarity was computed using correlation:

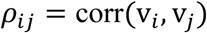

Bins with missing values were excluded pairwise. Similarities were assembled into square matrices and visualized as heatmaps.

### 4.5 Ks-Based Duplication Analyses

#### 4.5.1 Ks Filtering and Mixture Decomposition

Ks values associated with syntenic anchors were filtered to retain:

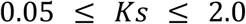

Filtered Ks values were modeled using Gaussian mixture models^43,83^ fitted by expectation-maximization.

Given Ks values {𝑥_1_, …, 𝑥*_n_*}, the model is:

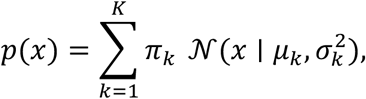

where 𝜋_𝑘_ denotes the mixture weight and 𝜇_𝑘_ and 𝜎_𝑘_ represent the mean and standard deviation of component 𝑘. Mixture fitting was performed using expectation-maximization, and model complexity was chosen conservatively to capture interpretable duplication signal rather than maximize statistical fit.

Model selection was performed using Bayesian Information Criterion:

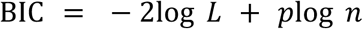

where 𝑝 = number of estimated parameters (for 1D GMM with 𝑘 components):

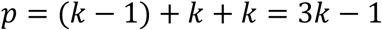

Components were sorted by increasing 𝜇*_k_* and treated strictly as relative age classes.

#### 4.5.2 Event-Stratified Synteny Density

Each anchor was assigned to a Ks component.

For each component 𝑒, reference chromosome 𝑐, and bin 𝑏:

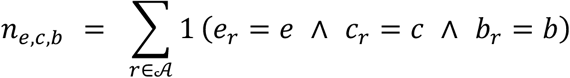

where:

- 𝑟 indexes syntenic anchor pairs
- 𝒜 is the set of all anchors retained for analysis
- 𝑘*_r_* is the Ks mixture component (event) assigned to anchor 𝑟
- 𝑐*_r_* is the reference chromosome to which anchor 𝑟maps
- 𝑏*_r_* is the reference-coordinate bin containing anchor 𝑟
- 1(⋅) is the indicator function (1 if true, 0 otherwise)

The per-chromosome probability normalization, i.e. each chromosome’s event-stratified density integrates to 1 across bins, is written as:

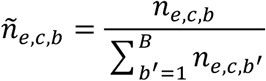

where:

- 𝑛*_e,c,b_* is the raw event-stratified anchor count (as previously defined)
- ñ*_e,c,b_* is the normalized density
- 𝐵 is the total number of bins on chromosome 𝑐
- the denominator sums only across bins on the same chromosome and event

As such, event-stratified anchor counts were normalized within each reference chromosome so that the summed density across bins equals one. These normalized densities were used for plotting age-stratified retention profiles.

#### 4.5.3 Ks Analysis and Conservative Duplication Inference Across Ericales

Ks distributions were analyzed to characterize the temporal structure of gene duplication within *Sarracenia purpurea* and to place these patterns within a comparative Ericales context. Duplicate gene pairs were obtained from self:self CoGe SynMap analyses^39–41^ for each species. Only gene pairs with defined chromosomal coordinates and valid Ks estimates were retained.

Ks values were filtered as in section 4.5.1 to reduce the influence of very recent noise at low divergence and extensive saturation at high divergence. Ks values outside this range were retained for visualization where informative but excluded from statistical modeling when dominated by multiple-hit effects. Filtered Ks values for each species were modeled using Gaussian mixture models as in section 4.5.1.

For all Ericales examined, the oldest and broadest Ks component was interpreted as retention of deeply diverged paralogs associated with the ancient core-eudicot *gamma* hexaploidy. This component was identified independently for each species and treated as a shared ancestral reference rather than evidence for recent duplication. Additional components at lower Ks were considered candidate post-*gamma* duplication signals only when they were distinct from the *gamma* component and not dominated by saturation or diffuse background signal.

Several Ericales taxa, including *Actinidia* and *Vaccinium*, exhibit complex Ks profiles with prominent mid- to low-Ks peaks consistent with additional lineage-specific whole-genome duplications reported for these genera. The presence of these recent WGDs obscures separation between *gamma*-derived and post-*gamma* duplication signal and complicates direct cross-taxon comparison of younger Ks modes. Accordingly, *Actinidia* and *Vaccinium* were retained for syntenic anchoring and contextual visualization but excluded from comparative inference of post-*gamma* duplication timing.

Comparative timing analyses therefore focused on *Sarracenia purpurea*, *Rhododendron*, and *Impatiens*, which lack strong evidence for additional recent polyploidy and exhibit simpler Ks landscapes. For these taxa, a single younger Ks component was modeled in addition to the *gamma* component. Component means were treated strictly as relative temporal markers rather than discrete event boundaries.

To place Ks components on a comparative time axis, Ks values were converted to divergence time using:

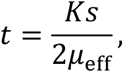

where 𝜇*_eff_* is an effective long-term synonymous substitution rate. For each species, 𝜇*_eff_* was calibrated independently by constraining the mean of the *gamma* component to 120-125 million years ago, corresponding to widely inferred estimates for the timing of core-eudicot *gamma* hexaploidy^45–47^. Younger component ages were then scaled relative to this species-specific calibration.

Because Ks-based age estimates are sensitive to mixture model behavior, saturation effects, and the quality of genome assemblies and annotations, inferred duplication times are reported as approximate and are interpreted comparatively rather than as precise chronological estimates. No assumption of homology among younger duplication components across taxa was imposed.

### 4.6 Repeat Analyses

#### 4.6.1 Repeat Divergence Landscape Analysis

Repeat divergence landscapes were generated from RepeatMasker annotations to provide a relative temporal summary of repeat accumulation in the *Sarracenia purpurea* genome. All RepeatMasker-annotated repeat fragments were treated as independent genomic intervals and analyzed without reference to gene models, syntenic structure, or chromosome identity.

For each annotated repeat fragment 𝑟, RepeatMasker reports a percent sequence divergence from the corresponding consensus element, reflecting accumulated substitutions since insertion relative to the library consensus. Let 𝑑_𝑟_ denote this reported divergence value (in percent).

Repeat fragments were binned into integer divergence classes spanning 0-50% divergence. Formally, each fragment 𝑟 was assigned to divergence bin:

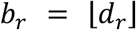

such that:

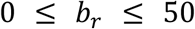

For each divergence bin 𝑏, total base-pair coverage was computed as the sum of fragment lengths:

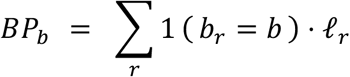

where ℓ*_r_* is the genomic length of fragment 𝑟. These base-pair totals were then normalized by the total assembled genome size 𝐺 to obtain bp-weighted divergence profiles:

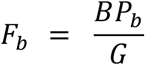

This procedure yields a genome-wide repeat divergence landscape in which each bin represents the fraction of the genome occupied by repeats with divergence values falling in that interval.

In addition to the genome-wide landscape, an LTR-restricted divergence landscape was constructed by applying the same binning and aggregation procedure to the subset of RepeatMasker fragments annotated as LTR-associated repeats. All normalization and bin definitions were identical between genome-wide and LTR-restricted analyses, enabling direct comparison of their divergence profiles.

For visualization, divergence landscape were plotted as histograms of 𝐹*_b_* across divergence bins, with smoothed curves overlaid to highlight modal structure. The smoothed curves were used solely for visualization and did not alter the underlying bin counts. The modal divergence bin for each landscape was identified as the bin 𝑏 with maximal 𝐹*_b_* and indicated graphically as a reference.

Further definitions:

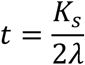

where 𝑡 is divergence time (in millions of years), 𝐾_s_ is the synonymous substitution rate, and 𝜆 is the effective long-term synonymous substitution rate.

Therefore:

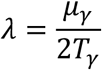

where 𝜇*_γ_* is the mean Ks value of the *gamma* component and 𝑇*_γ_* is the assumed timing of core-eudicot *gamma* hexaploidy (120-125 million years ago).

For each younger Ks component with mean 𝜇*_e_*, divergence time was then calculated as:

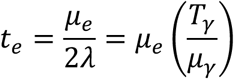

All inferred ages are interpreted comparatively rather than as precise chronological estimates.

Because RepeatMasker divergence values reflect sequence divergence from consensus rather than absolute insertion times, divergence landscapes were interpreted as relative summaries of repeat accumulation dynamics. Therefore, differences between genome-wide and LTR-restricted profiles were used to assess whether LTR retrotransposons contribute disproportionately to temporally structured components of repeat accumulation.

#### 4.6.2 Comparative Analysis of Ks-Based Duplication Signals and LTR Divergence Landscapes

To examine the temporal relationship between gene duplication and transposable-element accumulation within the *Sarracenia purpurea* genome, we integrated Ks-based duplication age estimates with RepeatMasker-derived divergence landscapes for LTR retrotransposons. Ks values were obtained from self-syntenic duplicate gene pairs, and in this case, filtered to the range 0.05 ≤ 𝐾𝑠 ≤ 2.0 to minimize the influence of saturation at high divergence.

Filtered Ks values were modeled using a two-component Gaussian mixture model, representing a younger lineage-specific duplication signal and a broader, older component corresponding to deeply diverged paralogs. The latter component is interpreted as reflecting retention from the ancient core-eudicot *gamma* hexaploidy event, while the younger component captures post-*gamma* duplication retained within the *Sarracenia* lineage. Mixture fitting was performed as in section 4.5.1.

LTR retrotransposon divergence landscapes were constructed by parsing RepeatMasker annotations to extract all LTR-associated repeat fragments. For each fragment, the reported percent divergence from its consensus sequence was binned into integer divergence classes, and base-pair coverage was summed per bin to generate bp-weighted divergence profiles.

To enable direct visual comparison between duplication and repeat accumulation, Ks values and LTR divergence bins were transformed to a common time axis using the relationship:

Where:

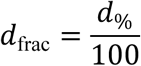

and then:

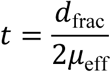

and 𝑑 denotes Ks or fractional divergence (see also section 4.5.3). An effective long-term substitution rate 𝜇*_eff_* was calibrated by constraining the mean of the older Ks component to 120-125 million years ago, corresponding to the widely inferred timing of the core-eudicot *gamma* hexaploidy event. All time estimates are therefore calibration-dependent and interpreted comparatively. Ks density curves and LTR divergence histograms were plotted on aligned time axes with independent y-axis normalization to emphasize relative temporal structure rather than absolute magnitude.

#### 4.6.3 Quantification of Transposable-Element Heterogeneity Across Pseudochromosomes

To quantify variation in transposable-element (TE) distribution across the *Sarracenia purpurea* genome, RepeatMasker annotations were summarized at the pseudochromosome scale using union base-pair coverage. All RepeatMasker-annotated intervals were parsed by pseudochromosome and merged prior to summation to avoid double-counting overlapping fragments. Total TE coverage was calculated as the number of base pairs covered by any RepeatMasker annotation (all TE classes combined) divided by the total length of each pseudochromosome.

LTR retrotransposon coverage was calculated analogously by restricting annotations to LTR-tagged elements, including Gypsy and Copia families, and computing union base-pair coverage relative to pseudochromosome length. Thus, heterogeneity was quantified separately for (i) overall TE occupancy and (ii) LTR-specific occupancy, enabling direct comparison between genome-wide repeat content and the dominant repeat class.

Inter-pseudochromosome variation was summarized using descriptive statistics across the 13 pseudochromosomes. For each metric, the range (Δ) was defined as the difference between the maximum and minimum coverage values, expressed in percentage points. The coefficient of variation (CV) was calculated as the standard deviation divided by the mean coverage and is reported as a percentage, providing a normalized measure of heterogeneity that accounts for differences in mean repeat abundance. These statistics were used to support statements regarding scaffold-level variation in repeat content and to contextualize the spatial distribution of LTR retrotransposons relative to overall TE composition.

Fragment counts per pseudochromosome were recorded as a measure of annotation density but were not used for coverage estimation. All reported percentages reflect union coverage and therefore represent conservative estimates of repeat occupancy.

### 4.7 Comparative Functional Genomics (Bins, Foregrounds, GOs)

#### 4.7.1 Gene Bin Construction Using Strict Intersection Logic

Protein-coding genes were first mapped to *Arabidopsis thaliana* ortholog identifiers, providing a common reference framework for cross-taxon comparison. Gene bins were then constructed using a strict intersection logic that encodes shared evolutionary presence patterns across a fixed set of taxa.

For a fixed set of taxa:

- For each *Arabidopsis* identifier, construct a binary presence-absence vector.
- Group identifiers with identical vectors.
- Each group defines a strict intersection bin.

This procedure partitions the full gene set into equivalence classes defined solely by shared taxonomic presence patterns. These bins form the atomic units for all downstream functional analyses.

Properties:

- Each gene belongs to exactly one bin.
- Bins are mutually exclusive.
- Bins are never modified downstream.

This binning step is intentionally conservative and is the only stage in the analysis where intersections across taxa are permitted. By fixing bin membership at this stage, all subsequent analyses inherit a stable, reproducible partitioning of genes that is independent of downstream biological hypotheses.

This is the only stage where intersections are used.

#### 4.7.2 Foreground Construction as Unions of Bins

All downstream gene sets used for functional analysis were defined strictly as unions of the pre-defined intersection bins described above. No additional filtering, subdivision, or intersection operations were applied once bins were established.

Explicitly:

- No genes were intersected across bins.
- No bins were intersected with other bins.
- Foregrounds are deterministic unions defined by bin membership.

This union-based strategy ensures that foregrounds remain internally consistent, reproducible, and traceable back to the original bin definitions. It also prevents inflation of statistical signal through repeated reuse or reshaping of overlapping gene sets.

This union-based foreground construction applies uniformly to:

- duplication-mode analyses
- subgenome compartment analyses
- lineage-specific gene sets

By enforcing this structure, functional comparisons across different biological questions (e.g., duplication mode versus lineage specificity) remain directly comparable and free from hidden dependencies.

#### 4.7.3 GO Enrichment

Gene Ontology (GO) enrichment analyses were performed on foregrounds defined as unions of bins, as described above. Enrichment testing was conducted using a standard contingency-table framework, with careful separation of genome-derived and expression-derived analyses.

For each GO term:

- Count foreground genes annotated to the term.
- Count background genes annotated to the term.
- Perform Fisher’s exact test.

The background consisted of all *Arabidopsis* identifiers eligible for bin assignment, ensuring that enrichment tests were conditioned on the same orthology universe used to define bins.

Genome-derived enrichments used Bonferroni correction:

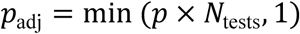

Expression-derived enrichments used Benjamini-Hochberg false discovery rate control, reflecting the higher degree of dependency expected among expression-based gene sets.

To avoid conflating broadly conserved Ericales functions with lineage-specific signal, GO terms enriched in *Actinidia*-containing foregrounds were identified separately and removed from *Sarracenia*-focused interpretations at the interpretation stage only. This filtering step was applied post hoc and did not alter the underlying enrichment calculations.

### 4.8 RNA-seq Processing and Expression Analyses

#### 4.8.1 RNA Sequencing and Sample Design

RNA sequencing was performed on three tissue types: roots, trap leaves, and flowers. Three biological replicates were collected for each tissue type, yielding a total of nine RNA-seq samples. Samples were designated as follows: SPR1-SPR3 (roots), SPL1-SPL3 (trap leaves), and SPF1-SPF3 (flowers). Flower samples were harvested after petal abscission to capture post-floral developmental expression states.

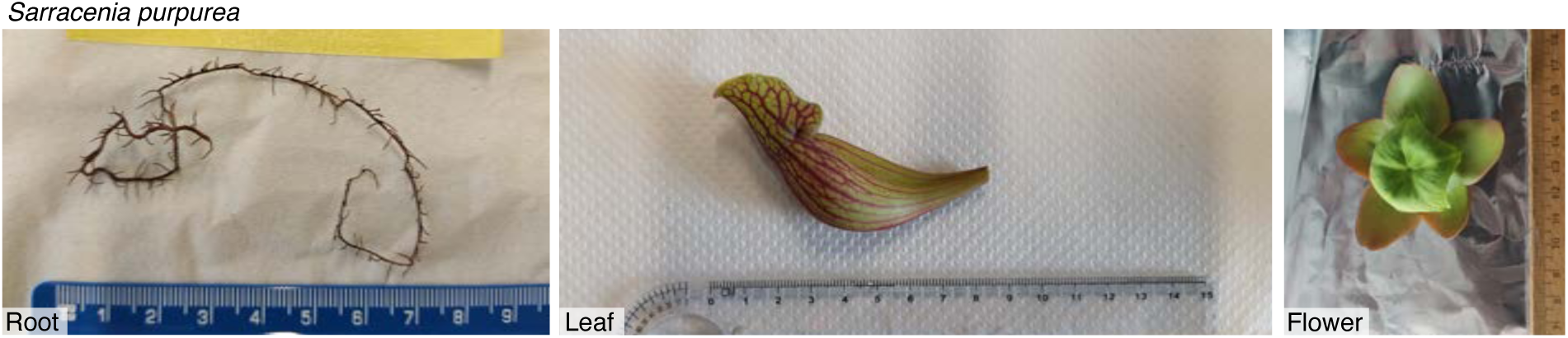

RNA-seq libraries were sequenced on an Illumina platform using paired-end reads. All RNA-seq data have been deposited under BioProject PRJDB15737. Detailed accession information for each sample is provided in section 4.9.

#### 4.8.2 RNA-seq Read Mapping and Transcriptome Assembly

Genome indexing for RNA-seq read alignment was performed using HISAT2 v2.2.1, incorporating splice-site and exon information derived from the final EVM-integrated gene annotation. Splice sites were extracted from the GFF3 annotation using *hisat2_extract_splice_sites.py*. Exon coordinates were obtained by converting the GFF3 file to GTF format using gffread v0.12.7, followed by *hisat2_extract_exons.py*.

The RagTag-scaffolded genome assembly was indexed using:

hisat2-build --ss Spur.ss --exon Spur.exons Spur.fasta --large-index -p 16 Spur_index

RNA-seq reads were aligned to the indexed genome using HISAT2 with parameters optimized for transcriptome assembly:

hisat2 --dta -p 16 -x Spur_index \

-1 SAMPLE_R1.fastq.gz -2 SAMPLE_R2.fastq.gz \

-S SAMPLE.sam

The --dta option was used to report alignments suitable for transcript assemblers. SAM files were converted to BAM format and sorted using SAMtools v1.16.1.

Transcript assembly and expression estimation were performed independently for each sample using StringTie v2.2.1:

stringtie SAMPLE.sorted.bam -G reference.gtf -e -B -o SAMPLE.gtf

The -e option restricted assembly to reference transcripts, and the -B option generated tabular output files required for downstream differential expression analyses.

#### 4.8.3 Differential Expression Analysis

Gene- and transcript-level count matrices were generated using *prepDE.py* following the StringTie workflow. This procedure produced a transcript count matrix (*transcript_count_matrix.csv*) summarizing expression counts for each annotated gene across all samples. These matrices were used for downstream differential expression analyses described in the Results.

#### 4.8.4 Expression Responsiveness of Syntenic Versus Tandem Duplicates

Expression responsiveness of duplicated genes in *Sarracenia purpurea* was compared between syntenically retained duplicates and tandem duplicates. Syntenic duplicates were defined as paralogous gene pairs retained within self-syntenic blocks identified through the Ericales anchoring framework, whereas tandem duplicates were defined as adjacent or near-adjacent paralogs identified using a CDS-based tandem detection pipeline. Genes classified as mixed or ambiguous with respect to duplication mode were excluded from this analysis.

Gene expression responsiveness was quantified using pitcher-associated transcriptomic datasets. For each gene, expression change was calculated as the absolute value of the log-transformed fold change across experimental contrasts, such that higher values reflect greater transcriptional responsiveness across pitcher-related conditions. This metric captures both up- and down-regulation while avoiding cancellation effects.

Distributions of expression responsiveness for syntenic and tandem gene sets were visualized using combined violin and box plots. Box plots indicate the median and interquartile range, while violins represent kernel density estimates of the full distribution. Differences between duplication classes were assessed using a two-sided Mann-Whitney U test, which does not assume normality. Effect size was quantified using Cliff’s delta (𝛿), which measures the probability that a randomly selected value from one group exceeds a randomly selected value from the other group, minus the reverse probability:

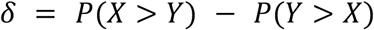

where 𝑋 and 𝑌 denote observations drawn from the two duplication classes. Sample sizes for each group were retained as observed and are reported directly on **Figure 14**. All statistical tests were conducted using gene-level values without subsampling.

#### 4.8.5 Functional Breadth Analysis Using GO Rarefaction

Functional breadth of syntenic and tandem gene sets was evaluated using Gene Ontology (GO) biological-process annotations under a rarefaction framework. GO annotations were derived from the curated genome annotation of *Sarracenia purpurea*, and genes lacking GO biological-process terms were excluded from this analysis.

For each duplication class, genes were repeatedly subsampled without replacement across a range of sampling depths. At each depth, two complementary measures of functional breadth were calculated. First, GO richness was defined as the number of unique GO biological-process terms represented in the subsample. Second, effective functional breadth (D1^84^) was calculated as the exponential of Shannon entropy (cf. ^81^):

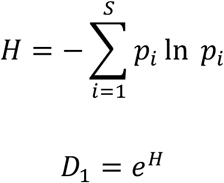

where 𝑝*_i_* is the proportional frequency of genes annotated to GO term 𝑖, and 𝑆 is the total number of GO terms observed in the subsample. This metric captures both the diversity and evenness of functional annotations.

For each sampling depth, mean values and confidence envelopes were estimated from the distribution of subsampled values across iterations. Because rarefaction curves represent non-independent function-valued data, no formal hypothesis testing was applied across entire curves. Instead, rarefaction was used as a descriptive framework to visualize differences in functional breadth between duplication classes while controlling for gene set size.

### 4.9 RNA-seq Samples and Accession Metadata

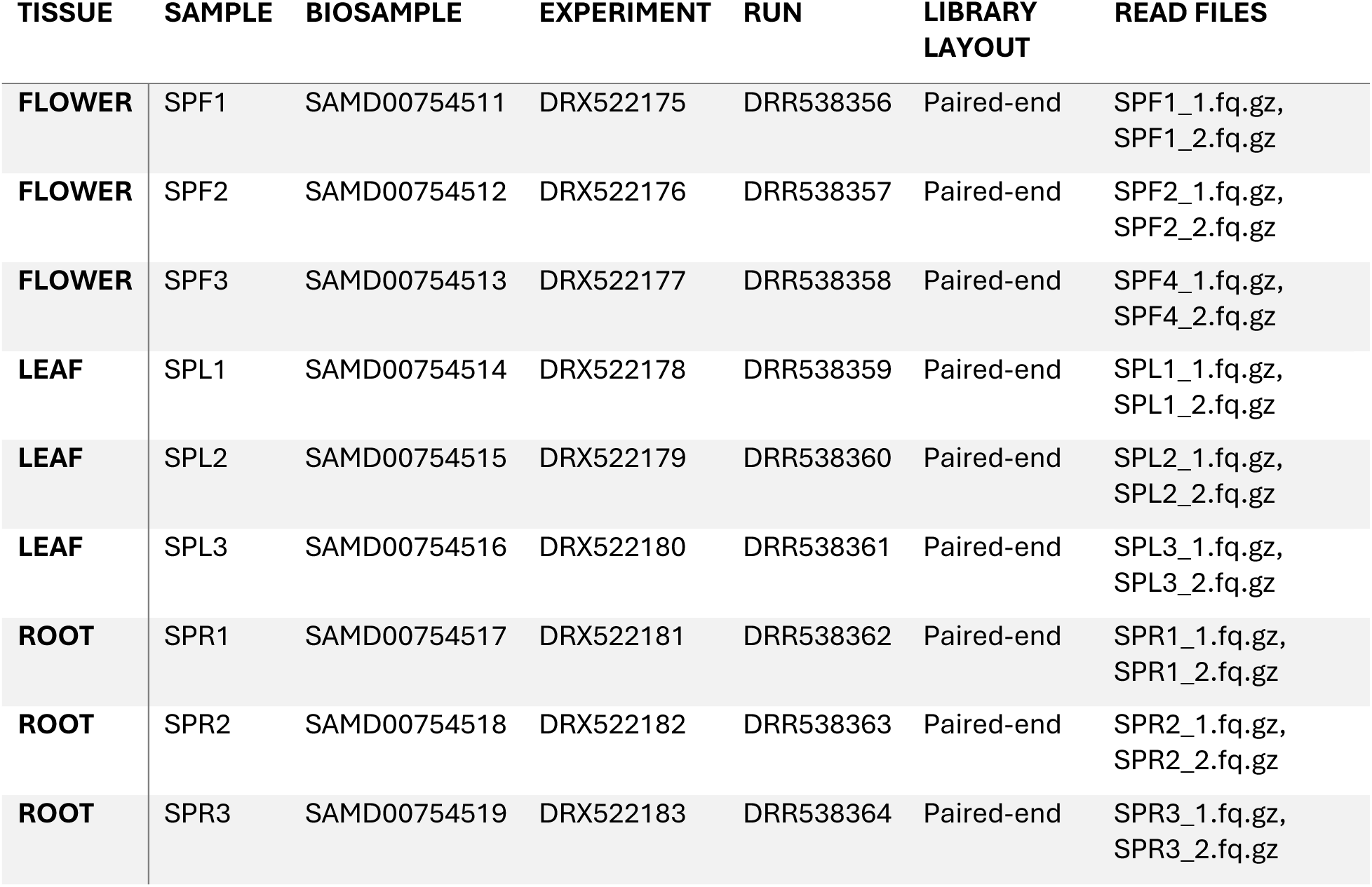

| TISSUE | SAMPLE | BIOSAMPLE | EXPERIMENT | RUN | LIBRARY LAYOUT | READ FILES |
| --- | --- | --- | --- | --- | --- | --- |
| **FLOWER** | SPF1 | SAMD00754511 | DRX522175 | DRR538356 | Paired-end | SPF1_1.fq.gz,<br>SPF1_2.fq.gz |
| **FLOWER** | SPF2 | SAMD00754512 | DRX522176 | DRR538357 | Paired-end | SPF2_1.fq.gz,<br>SPF2_2.fq.gz |
| **FLOWER** | SPF3 | SAMD00754513 | DRX522177 | DRR538358 | Paired-end | SPF4_1.fq.gz,<br>SPF4_2.fq.gz |
| **LEAF** | SPL1 | SAMD00754514 | DRX522178 | DRR538359 | Paired-end | SPL1_1.fq.gz,<br>SPL1_2.fq.gz |
| **LEAF** | SPL2 | SAMD00754515 | DRX522179 | DRR538360 | Paired-end | SPL2_1.fq.gz,<br>SPL2_2.fq.gz |
| **LEAF** | SPL3 | SAMD00754516 | DRX522180 | DRR538361 | Paired-end | SPL3_1.fq.gz,<br>SPL3_2.fq.gz |
| **ROOT** | SPR1 | SAMD00754517 | DRX522181 | DRR538362 | Paired-end | SPR1_1.fq.gz,<br>SPR1_2.fq.gz |
| **ROOT** | SPR2 | SAMD00754518 | DRX522182 | DRR538363 | Paired-end | SPR2_1.fq.gz,<br>SPR2_2.fq.gz |
| **ROOT** | SPR3 | SAMD00754519 | DRX522183 | DRR538364 | Paired-end | SPR3_1.fq.gz,<br>SPR3_2.fq.gz |

## 5. Data Availability

The genome assembly and annotation of *Sarracenia purpurea* is publicly available at https://genomevolution.org/coge/GenomeInfo.pl?gid=69349. RNA-seq data are available at NCBI BioProject PRJDB15737. All other data can be requested from the corresponding author (Victor A. Albert) and will also be made available upon formal journal publication.

## 6. Acknowledgments

We acknowledge the following sources for funding: the U.S. National Science Foundation grants 2030871 and 2139311 (to VAA and CL, respectively), the Sofja Kovalevskaja Program of the Alexander von Humboldt Foundation (to KF), and JSPS KAKENHI 23K20050 (to KF).

## 7. Author Contributions

VAA conceived the study; VAA led the research with assistance from JK; VAA, JK, MR, and CL provided materials; MR sequenced Oxford Nanopore reads; KF and MF provided RNA-seq data; CP, JK, and NP assembled and annotated the genome; CP decontaminated and reference-scaffolded the assembly; CP analyzed RNA-seq data; VAA, JK, CP, NP, and JM performed additional downstream bioinformatic analyses; VAA, CP, and JM produced graphics; VAA wrote the manuscript, and all authors approved the final version.

## 9. Supplementary Tables

**Table S1.** Summary of gene and transcript structural features in the *Sarracenia purpurea* genome. Key annotation metrics for the primary gene set, including counts and length distributions for genes, transcripts, coding sequences, exons, and introns. Values summarize overall genome architecture, highlighting a moderately exon-rich gene space, substantial intron-length variation, and a genome-wide gene density of about 8 genes per megabase across the 3.52 Gb assembly. Each gene locus was represented by a single representative transcript model in the final annotation.

| METRIC | VALUE |
| --- | --- |
| NUMBER OF GENE LOCI | 28,835 |
| NUMBER OF TRANSCRIPTS | 28,835 |
| TOTAL CDS FEATURES | 132,001 |
| MEAN EXON LENGTH (BP) | 263.12 |
| MEDIAN EXON LENGTH (BP) | 139 |
| MEAN GENE LENGTH (BP) | 9,315.68 |
| MEDIAN GENE LENGTH (BP) | 3,112 |
| MEAN TRANSCRIPT LENGTH (BP) | 9,315.68 |
| MEDIAN TRANSCRIPT LENGTH (BP) | 3,112 |
| MEAN EXON COUNT PER GENE | 4.58 |
| MEDIAN EXON COUNT PER GENE | 3 |
| NUMBER OF SINGLE-EXON GENES | 7,179 |
| MEAN INTRON LENGTH (BP) | 2,267.08 |
| MEDIAN INTRON LENGTH (BP) | 558 |
| GENOME-WIDE GENE DENSITY (GENES/MB) | 8.18 |
| TOTAL ASSEMBLY LENGTH (BP) | 3,523,999,544 |

**Table S2.**
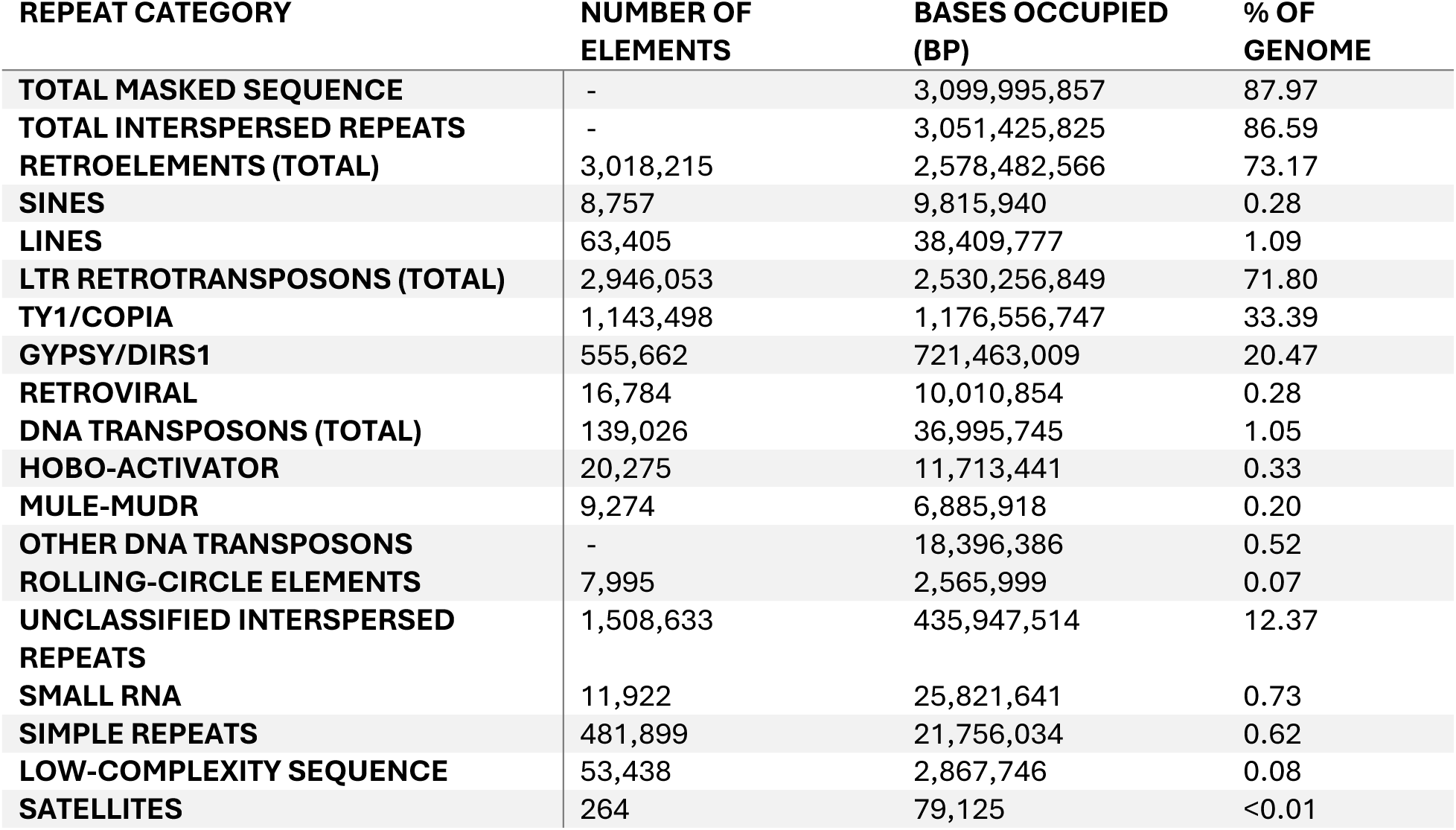
Genome-wide distribution of transposable element classes and other masked repeat categories in the *Sarracenia purpurea* assembly. Summary of RepeatMasker annotations showing the proportional contributions of major TE groups, including LTR retrotransposons, LINEs, SINEs, DNA transposons, rolling-circle elements, and unclassified repeats. The table also reports ancillary repeat categories such as small RNAs, simple repeats, and low-complexity regions, providing an overview of the repeat landscape that shapes genome size, structure, and gene-density variation in *Sarracenia purpurea*.

**Table S3.** *Sarracenia*-specific functional innovation identified by comparative foreground analyses. This table summarizes biological-process categories and representative genes associated with *Sarracenia*-specific and *Sarracenia*-enriched innovation foregrounds identified through comparative gene-set analyses across carnivorous plants and close Ericales relatives. Functional categories shown derive from gene sets present in Sarracenia but absent from, or explicitly filtered relative to, *Actinidia*, as well as from broader carnivore-union foregrounds. Representative *Arabidopsis* gene identifiers are provided to facilitate functional interpretation, with corresponding Sarracenia homologs listed where available. Duplication-mode annotations indicate whether genes occur among tandem-derived and/or syntenically retained loci; no enrichment, dominance, or causal relationship is implied. Rows reflect curated extractions from previously generated innovation and GO summary tables, and no additional statistical analyses were performed.

| FUNCTIONAL CATEGORY | REPRESENTATIVE GO BIOLOGICAL PROCESS | FOREGROUND CLASS | ACTINIDIA FILTER | ARABIDOPSIS GENE(S) | DUPLICATION MODE(S) |
| --- | --- | --- | --- | --- | --- |
| **DETOXIFICATION / XENOBIOTIC TRANSPORT** | Response to toxic substance (GO:0009636) | Carnivore-union | Shared with *Actinidia* | AT1G01580 (*PEN3*-like ABC transporter) | Tandem, syntenic |
| **GLUTATHIONE-BASED REDOX REGULATION** | Glutathione metabolic process (GO:0006749) | Carnivore-union | Shared with *Actinidia* | AT2G16390 (*GSTF* family) | Tandem |
| **ANTIFUNGAL DEFENSE** | Defense response to fungus (GO:0050832) | *Sarracenia*-only | Absent from *Actinidia* | AT2G26560 | Tandem |
| **WOUND / STRESS RESPONSE** | Response to wounding (GO:0009611) | *Sarracenia*-only | Absent from *Actinidia* | AT1G24140 | Tandem, syntenic |
| **CUTICLE BIOSYNTHESIS / REMODELING** | Cuticle development (GO:0042335) | Carnivore-union | Shared with *Actinidia* | *CYP86*- and *LACS2*-associated genes | Tandem |
| **ETHYLENE-ACTIVATED SIGNALING** | Ethylene-activated signaling pathway (GO:0009873) | *Sarracenia*-only | Absent from *Actinidia* | AT-annotated regulators | Mixed |
| **CELL DIFFERENTIATION** | Cell differentiation (GO:0030154) | *Sarracenia*-only | Absent from *Actinidia* | AT-annotated regulators | Mixed |
| **ROOT-HAIR CELL DIFFERENTIATION** | Root hair cell differentiation (GO:0048767) | *Sarracenia*-only | Absent from *Actinidia* | AT-annotated regulators | Mixed |
| **MITOCHONDRIAL RNA MODIFICATION** | Mitochondrial mRNA modification (GO:0042788) | *Sarracenia*-only | Absent from *Actinidia* | AT-annotated factors | Mixed |
| **CHROMATIN-LEVEL REGULATION** | DNA demethylation (GO:0080111) | *Sarracenia*-only | Absent from *Actinidia* | AT2G36490 (*DML1*) | Tandem |

**Tables SA-SI.** See external Zip file: Tables_SA-SI.zip

