## Supplementary Tables SA-SI for "Genome evolution and convergent innovation in the carnivorous plant *Sarracenia purpurea*": README.txt

PRIMARY TABLE PACKAGE


CONTENTS
--------

Table_SA_SarraceniaOnly_Innovation_Genes_Summary.csv
  Purpose: Sarracenia-only innovation gene summary with duplication-mode flags

Table_SB_Sarracenia_Nepenthes_GO_Union_Enrichment.csv
  Purpose: Sarracenia–Nepenthes GO-union enrichment results with genes per lineage

Table_SC_PanCarnivore_GO_Modules.csv

Table_SD_CarnivoreOnly_GeneFamily_Categories.csv
  Purpose: Carnivore-only gene families categorized by functional class across taxa

Table_SE_MultiTaxon_GO_Bin_Summary.csv
  Purpose: Multi-taxon GO bin summary with Sarracenia-specific filter flags

Table_SF_Sarracenia_Specific_GO_Mapping_Per_ATGene.csv
  Purpose: Per-gene mapping of Sarracenia-specific GO IDs and names

Table_SG_SarraceniaOnly_Genes_with_DEG_Support.csv
  Purpose: Sarracenia-only gene list with duplication-mode and DEG support flags

Table_SH_Expression_Support_for_Key_Families.csv
  Purpose: DEG status and innovation-class support for key families (PEN3, DML1, etc.)

Table_SI_Sarracenia_Gene_to_Arabidopsis_Orthology_and_Context.csv
  Purpose: Sarracenia genes with Arabidopsis mapping and Vitis-anchored context (for gene-level callouts)
